# Synchrony dynamics underlie irregular neocortical spiking

**DOI:** 10.1101/2024.10.15.618398

**Authors:** Jagruti J. Pattadkal, Ronan T O’Shea, David Hansel, Thibaud Taillefumier, Darrin Brager, Nicholas J. Priebe

**Affiliations:** The University of Texas at Austin; CNRS; University of Nevada at Las Vegas

## Abstract

Cortical neurons are characterized by their variable spiking patterns. We challenge prevalent theories for the origin of spiking variability. We examine the specific hypothesis that cortical synchrony drives spiking variability *in vivo*. Using dynamic clamp, we demonstrate that intrinsic neuronal properties do not contribute substantially to spiking variability, but rather spiking variability emerges from weakly synchronous network drive. With large-scale electrophysiology we quantify the degree of synchrony and its time scale in cortical networks *in vivo*. We demonstrate that physiological levels of synchrony are sufficient to generate irregular responses found *in vivo*. Further, this synchrony shifts over timescales ranging from 25 to 200 ms, depending on the presence of external sensory input. Such shifts occur when the network moves from spontaneous to driven modes, leading naturally to a decline in response variability as observed across cortical areas. Finally, while individual neurons exhibit reliable responses to physiological drive, different neurons respond in a distinct fashion according to their intrinsic properties, contributing to stable synchrony across the neural network.

## Introduction

The variability of neurons has long stood as a central and classical feature of cortical neurons. In sensory areas like visual cortex, distinct patterns of action potentials are observed in response to repeated presentations of the same sensory stimulus (1-3). Similarly, neurons in premotor and motor cortices variably respond when animals are instructed to execute the same action (4). Spiking variability could arise from noisy synaptic inputs that converge to individual neurons (5-8), as well as from inherently intrinsic stochastic cellular mechanisms (9-11). *In vitro* recordings have revealed that neurons respond unreliably to steady input, presumably due to stochastic cellular processes but that large input fluctuations can overcome this stochasticity and lead to reliable responses, though the physiological relevance of either of these input regimes is unclear (12).

Previous theoretical studies argued that excitatory drive generally fails to exhibit variable responses, as neurons are thought to integrate the input from large numbers of asynchronously spiking neurons (6, 13-16). Variable responses may emerge, however, from the convergence of strong excitatory and inhibitory asynchronous inputs. In this condition, the mean drive from these sources cancels out, but their variability remains (5, 6, 17, 18). The aim of the present study was to determine experimentally the physiological conditions that generate variable spiking in cortical neurons.

### Spiking variability is not due to intrinsic neuronal properties but due to network dynamics

We first examined the relative contributions of synaptic and intrinsic cellular sources of noise by making whole-cell conductance dynamic clamp recordings from pyramidal neurons in mouse and marmoset cortical slices. To emulate *physiological* input conditions, we injected excitatory and inhibitory conductances which we previously recorded *in vivo* in visual cortex with and without visual stimulation (19). We initially adjusted the conductance gains to evoked between 3 and 21 spikes/s (mean = 11.5 spikes/s, range 3.7-21.3). Physiological drive evoked highly variable spiking patterns, characterized by an ISI coefficient of variation (CV) near 1 (CV of ISI: 1.2 ± 0.2 s.d., 12 cells, input source 1, 0.8 ± 0.2 s.d., 11 cells, input source 2) (Fig. 1A, Sup. Fig 1-1) (5, 13, 14).

**Figure 1:**
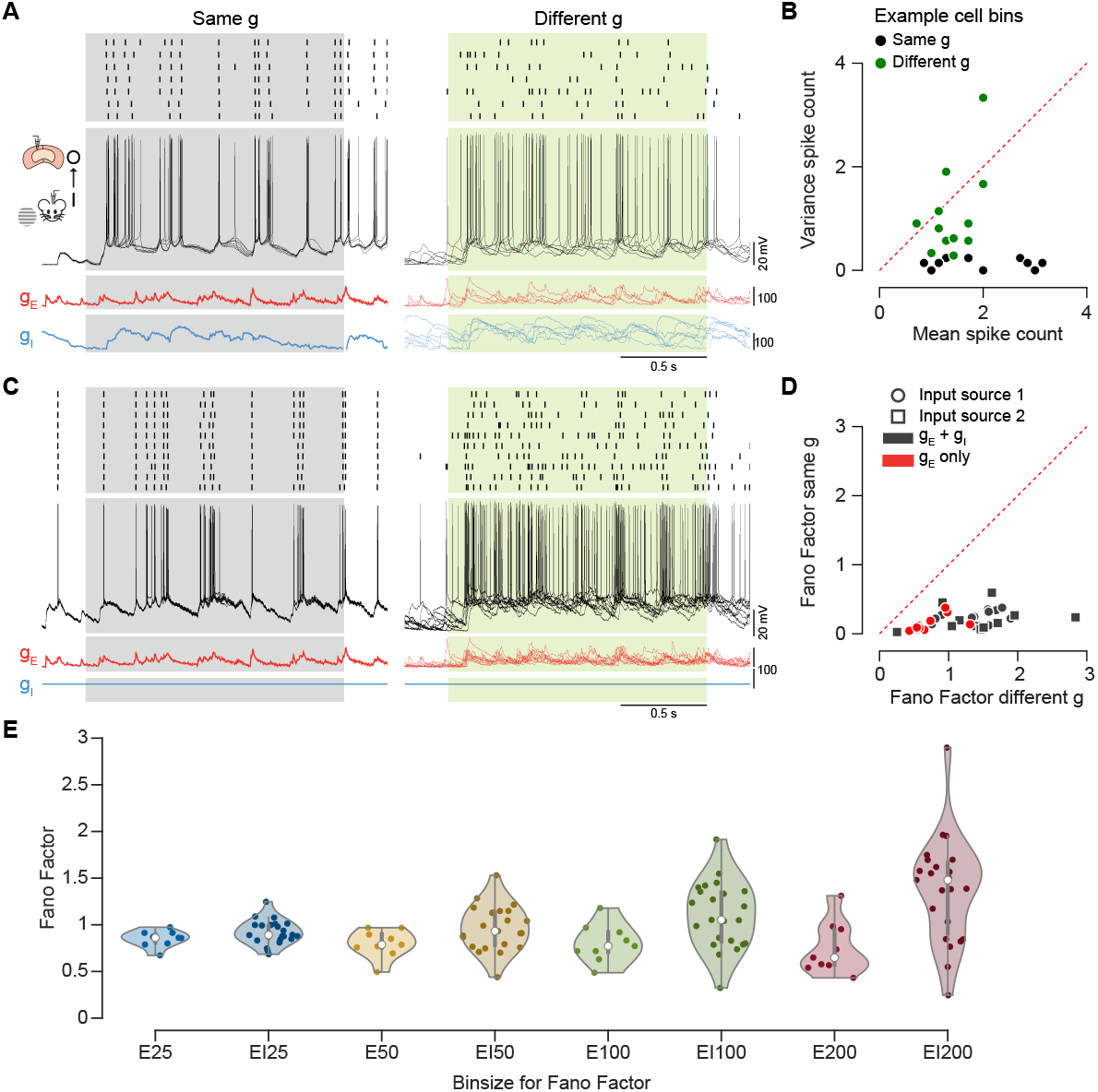
The reliability and variability of cortical neurons. A. Dynamic clamp was used *in vitro* to present excitatory (red) and inhibitory (blue) conductances recorded *in vivo*. Membrane potential and spiking responses are shown for repeating the same conductances (left), or for different combinations of excitatory and inhibitory conductance (right) in an example cell. B. The mean and variance of the spike count was measured in 200 ms bins for the same conductance condition (black circles) or the different conductance condition (green circles). Mean and variance of each bin for cell in A are shown. C. As in A, but the inhibitory conductance was set to 0. D. Fano factor across neurons for the same and different conductance conditions (n = 12 cells for input source 1, 11 cells for input source 2, 9 cells for excitatory conductance only). E. The Fano Factors for the different conductance conditions are shown of different time bins.

If intrinsic cellular processes contribute significantly to spiking variability, we expect to observe different spiking patterns across repeated presentations of the *same* physiological conductance drive. Contrary to this expectation, we found that repeated injections of the same excitatory and inhibitory conductances yielded precise patterns of action potentials with little difference in both marmoset and mouse neocortical neurons *in vitro* (Fig. 1A). To quantify this observation we computed the spike-count Fano factor, defined as the ratio of the variance and the mean of the spike count in a fixed time bin (Fig 1B). We found that repeatedly driving neurons *in vitro* with the same *in vivo*-measured conductances yielded low Fano factors (Fano factor: 0.3 ± 0.1 s.d. for input source 1, 12 cells, 0.3 ± 0.4 s.d. for input source 2, 11 cells) indicating a low degree of trial-to-trial spike count variability (Fig. 1B,D). In contrast, the Fano factor is near 1 for visual cortical responses to visual stimuli *in vivo* (20-22). Because neuronal responses are highly consistent across repeated injections of the same physiological drive, we conclude that intrinsic variability is unlikely to be a major source of cortical response variability and we term these responses ‘quasi-deterministic’ (23).

As intrinsic processes contribute little to response variability across repeats, the spiking variability measured *in vivo* across multiple presentations of the same visual stimulus must stem from variability in synaptic drive (Fig. 1A). To confirm this, we measured synaptic conductances evoked *in vivo* by repeated presentations of the same visual stimulus. We then injected these conductances *in vitro* to determine whether the cross-trial synaptic input variability could account for the variability of neuronal discharges. In contrast to the repeatedly injecting the same conductance (Fig. 1A, left), these cross-trial conductances resulted in highly variable trial to trial responses for physiological mean spiking rates (Fig. 1A, right). We found that the Fano factor for neurons in this condition is larger than 1 (Fano factor: 1.7 ± 0.8 s.d. for input source 1, 12 cells, 2.8 ± 1.3 s.d. for input source 2, 11 cells) in both marmosets and mice (Fig. 1B, Sup. fig. 1-1), a result that persists when varying the bin size used to count spikes (Sup Fig. 1-1). As expected, emulating trial variability in the synaptic drive allowed us to recapitulate *in vivo* spiking variability *in vitro*.

One caveat of our approach is the uncertainty about how to set the gain for the injected conductances. Measuring conductances *in vivo* requires blockade of voltage-gated channels, such as those responsible for the generation of action potentials (19, 24-27). Because of this alteration, our conductance measurements are only scaled versions of the true conductances by an unknown gain parameter. Given this uncertainty, we systematically varied the gain of both excitatory and inhibitory inputs and measured the resulting changes in firing rate and spiking variability. Increasing excitatory and inhibitory conductance gain equivalently led to a consistent change in firing rate but little change in the Fano factor (Sup. Fig. 2). Therefore, and perhaps surprisingly, although the overall firing rate changes, our measurements of spiking variability did not depend on the strength of the presented conductances.

Taken together, these results demonstrate that cortical spiking variability *in vivo i*s not due to intrinsic neuronal properties but instead emerges from the network dynamics, which generate variations in the synaptic drive to individual neurons for the same visual stimulus (3).

### Excitatory conductances alone produce Poisson-like spiking

Previous spiking network modeling studies have demonstrated that excitatory drive alone, generated by a large pool of neurons firing independently, leads to regular, clock-like spiking, with a low CV of ISI and a low Fano factor and (13). To test whether this is also the case for physiological drive, we presented excitatory conductances alone and measured the resulting Fano factor under two conditions: (1) for repeated presentations of the same *in vivo*-measured conductance traces (Fig. 1C, left), and (2) for different *in vivo*-measured conductance traces obtained in response to the same visual stimulus (Fig. 1C, right). With excitation alone, we found that neurons responded consistently in condition 1 (Fano factor: 0.1 ± 0.2 s.d., 9 cells) and variably in condition 2 (Fano factor: 0.8± 0.5 s.d., 9 cells), with degrees of consistency and variability similar to the responses observed with inhibition. Therefore *in vivo* excitatory conductances alone are sufficient to generate Poisson-like spiking variability *in vitro*.

### Statistics of *in vivo* conductances are inconsistent with asynchronous spiking

Our observation that excitatory drive alone is sufficient to generate spiking variability appears to be at odds with theoretical predictions (13, 14). These predictions stemmed from two critical assumptions. First, that presynaptic neurons are spiking independently as suggested by paired extracellular recordings in which correlations between neurons are weak (28). Second, anatomical studies show that cortical neurons receive synaptic contacts from a large pool of upstream neurons (29-32), thus the number of synaptic inputs is also likely to be large. By the law of large numbers, these two assumptions imply that cortical excitation alone should be a steady drive with very weak fluctuations, causing cortical neurons to spike regularly, contrary to our experimental observations. This apparent inconsistency can be resolved when estimating the amount of synchrony required to account for the fluctuations observed in our in vivo recordings.

To show that, let us assume that the activation of each connection is governed by a Poisson process so that the resulting conductances can be modeled as shot-noise traces. For simplicity, we study the impact of synchrony on the aggregate conductance over all the connection contributions. The temporal mean of the aggregate conductance is independent of the degree of synchrony and is given by *μ*_*g*_ = *AKrτ*_*s*_, where *A* is the typical size of a single EPSP, *K* is the number of inputs, *r* is the mean individual synaptic rate, and *τ*_*s*_ is the synaptic time constant. In contrast the temporal fluctuations of the aggregate conductance depend on synchrony. For independent firing, their standard deviation scales as 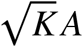. On another hand, for perfect synchrony the standard deviation scales as *KA*, as if the conductance changes resulted from the action of a single synapse of size *KA*. More generally, one can determine how the conductance variance depends on the pair-wise spiking correlation, *ρ* (see methods, (33)). Following on these observations and assuming a known synaptic time constant *τ*_*s*_, one can use the measured *μ*_*g*_ and *σ*_*g*_ to form the gain-independent quantity:

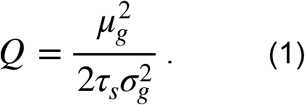

In the absence of synchrony, *i*.*e*., *ρ* = 0, *Q* is a measure of the aggregate input rate to a cell: *Kr*. Given the aggregate input rate we can then estimate the number of connections and compare those to the number of anatomical connections (Fig. 2A). Applying the same analysis to our *in vivo* conductance measurements yielded aggregate input rates that are remarkably low for both excitation and inhibition during visual stimulation (Excitation: median = 69, range = 10-173 Hz, Inhibition: median = 31, range = 1-235 Hz), and during spontaneous activity (Excitation: median = 18, range = 5-21 Hz, Inhibition: median = 30, range = 10-59 Hz, Fig. 2B,C). Our estimates of the aggregate input rate are much lower than the large number of presynaptic contacts observed anatomically (between 1000 and10000) and the firing rates (1-100) observed physiologically (30, 34-39).

**Figure 2:**
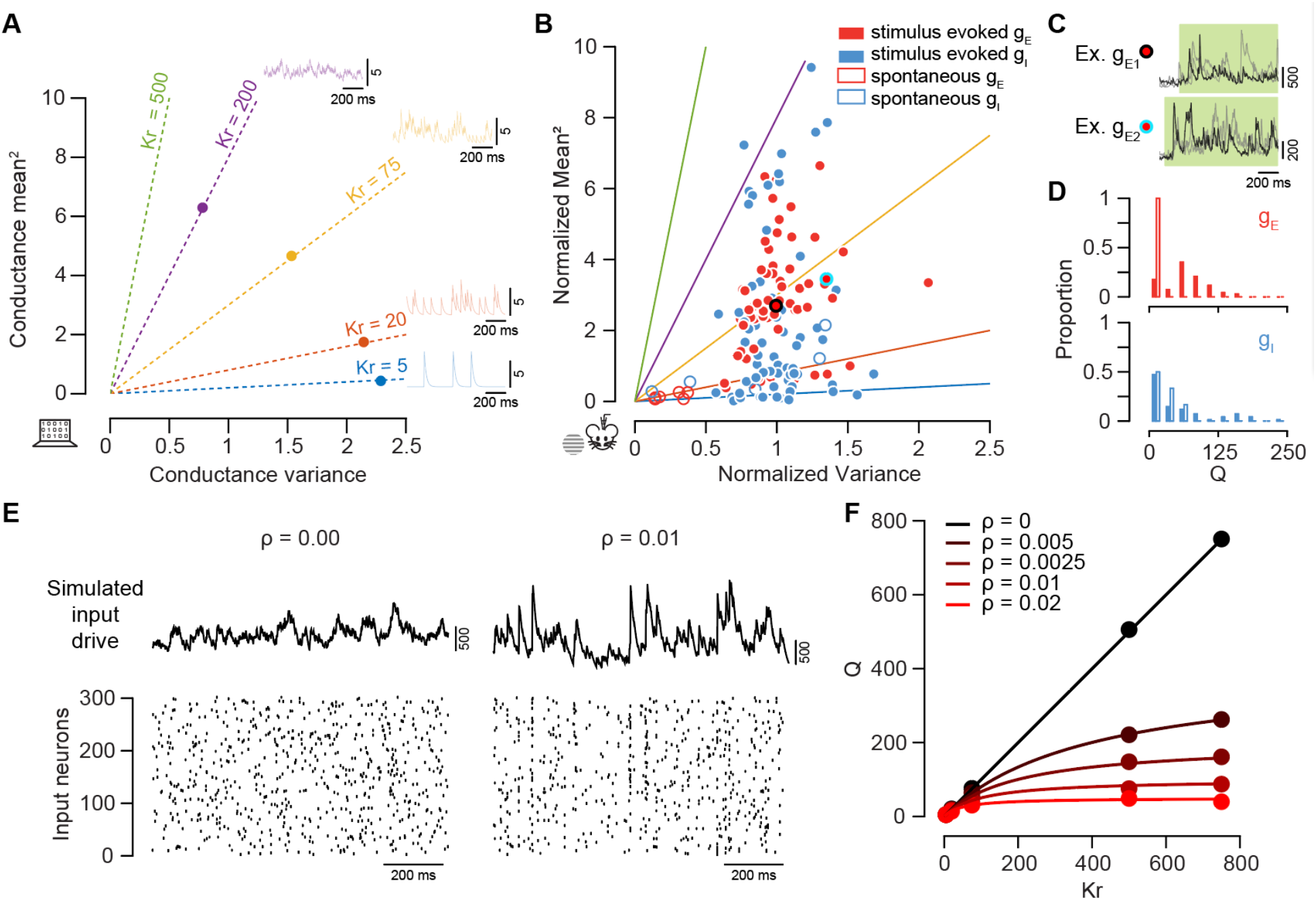
Estimates of the number of inputs onto cortical neurons. A. A model demonstrating how the slope of the relationship between the aggregate input variance and mean-squared relates to the number of inputs a neuron receives per second. One-second-long examples of conductance traces are shown above, with color indicating the rate of asynchronous inputs. B. The relationship between input variance and mean-squared among in vivo conductance measurements. Each point represents conductance for a cell for one stimulus condition, 6 cells are used here. C. Example repeats for two of the points shown in B. D. Histograms of the input rate, R, for both excitation and inhibition. E. Example spike rasters and net resulting drive from a population of neurons with pairwise correlations = 0 (left) or 0.01 (right). F. The relationship between input rate Kr in model neuronal populations relative to the Q estimated using the asynchronous assumption. The rate, r, is set to 1 spks/s. Color indicates different pair-wise correlations, symbols are based on simulated models and the solid line indicates the analytical prediction (see Methods).

Although estimates vary between reports (28, 40), *in vivo* extracellular recordings have revealed ‘weak’ spiking correlations among pairs of neurons. We hypothesized that these weak correlations might have an impact that cannot be neglected when estimating *Kr* (41). The inclusion of input synchrony causes the measurable quantity *Q* to underestimate the true values of *Kr* by a factor 1/(1 + *ρ*(*K* − 1)) (see Methods). Moreover, these underestimates saturate in the limit of large input numbers with 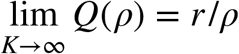, consistent with the fact that increasing level of spiking correlation leads to lower estimates of *Q*(*ρ*). We next simulated drives with varying number of inputs and varying degree of spiking correlation, which replicated our analytical derivations (Fig. 2E). These analyses and simulations reveal that even a modest level of spiking correlation (28, 40), *e*.*g*., *ρ* = 0.01, yields estimates of *Kr* that are consistent with realistic values for input numbers and synaptic rates (Fig. 2F).

### *In vivo* population measurements are consistent with cortical synchrony

The above results suggest that synchrony in the spiking activity of pre-synaptic neurons which is as weak as *ρ* = 0.01 could account for aggregate conductance statistics similar to those we observed experimentally. To determine whether such spiking correlations are present *in vivo*, we made large-scale measurements of cortical responses using neuropixels in marmosets and mice. Our measurements revealed clear bouts of synchronized responses on individual trials, which significantly deviate from the mean responses across trials. It was, however, difficult to quantify the degree of synchrony of these responses via direct estimates of pairwise correlations because of the presence of population and temporal heterogeneities.

To address these limitations, we use a population-level synchrony metric *χ*, which we modified from prior work (42, 43) and defined as

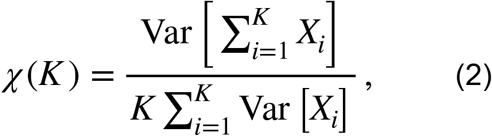

This *χ*-metric has been proposed as a general measure of synchrony for large population recordings (42, 43). In our case, *X*_*i*_ represents the varying spike counts of upstream neuron *i*, with ≤ *i* ≤ *K*. To gain intuition about the -metric, let us consider an homogeneous population of inputs which also acts independently across time bins. In such a case, by independence across neurons, we have Cov(*X*_*i*_, *X*_*j*_) = 0, and the population variance (numerator in (2)) computed over distinct time bins behaves additively over the neurons. As a result, we have *χ* (*K*) = 1/*K*. By contrast, in the presence of synchrony, neurons tend to activate together, which leads to positive spike-count correlations: Cov(*X*_*i*_, *X*_*j*_) > 0. Due to the inclusion of *K*(*K* – 1) additional cross terms Cov(*X*_*i*_, *X*_*j*_), ≤ *i* ≠ *j* ≤ *K*, the population variance (numerator in (3)) no longer behaves additively. As a result, we have

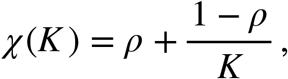

where *ρ* = Cov [*X*_*i*_, *X*_*j*_]/ Var[*X*_*i*_] can be rigorously interpreted as the pairwise spiking-correlation coefficient. One can therefore estimate input synchrony by examining the linear dependence of *χ* as a function of 1/*K* and extract the pairwise correlation coefficient,*ρ*, as the -intercept. The above arguments generalize, albeit with some caveats, to heterogeneous neuronal populations with time-varying firing rates (see Appendix). In particular, compared with classical use of the *χ*-metric, this generalization computes variance estimates across trials for each time bin, and then average these variance estimates across time bins, thereby allowing for variable population firing rates across neurons and time bins.

Applying the *χ*-metric to our measurements in the visual cortex of awake marmosets and mice revealed clear evidence for synchronous spiking activity (Fig. 3A, Sup. Fig. 3). Specifically, we found that Eq. 2 provides an excellent linear fit to the data and that extrapolation to 1/*K* → 0 consistently yields a *y*-intercept which is significantly different from zero. Since this intercepts measures the spiking synchrony at the population-level, this indicates that the spiking activity is significantly correlated across neurons (Fig. 3B,C,D). We found that the degree of spiking correlations varied between recordings, with mean *ρ* of 0.014 ± 0.014 s.d. (n = 13 marmoset cortical populations) during visual stimulation and mean *ρ* of 0.027 ± 0.025 s.d. (n = 10 marmoset cortical populations) during spontaneous activity, when measured for 200ms-long bins in awake marmosets (Fig. 3E, Sup Fig 3 for mice). The synchronous responses are a feature of not just the cortical populations, but also of LGN population (Sup Fig 3B) which sends input to the visual cortex (44). The degree of correlation also increased upon selecting subpopulation of visually responsive cells (Sup Fig 4). To assess the significance of our correlation estimates *ρ* we performed our *χ* -metric analysis on surrogate data whereby spikes emitted by the same neuron and within the same time bin are shuffled across trials. Such shuffling erases all spiking correlations while preserving population and temporal rate heterogeneities. Consistently, we found that the *χ*-metric analysis of these surrogate data yielded near-zero correlation estimates that were much smaller than the estimates we obtained on real data, confirming their significance (Fig. 3C,D,E, Sup Fig 3).

**Figure 3:**
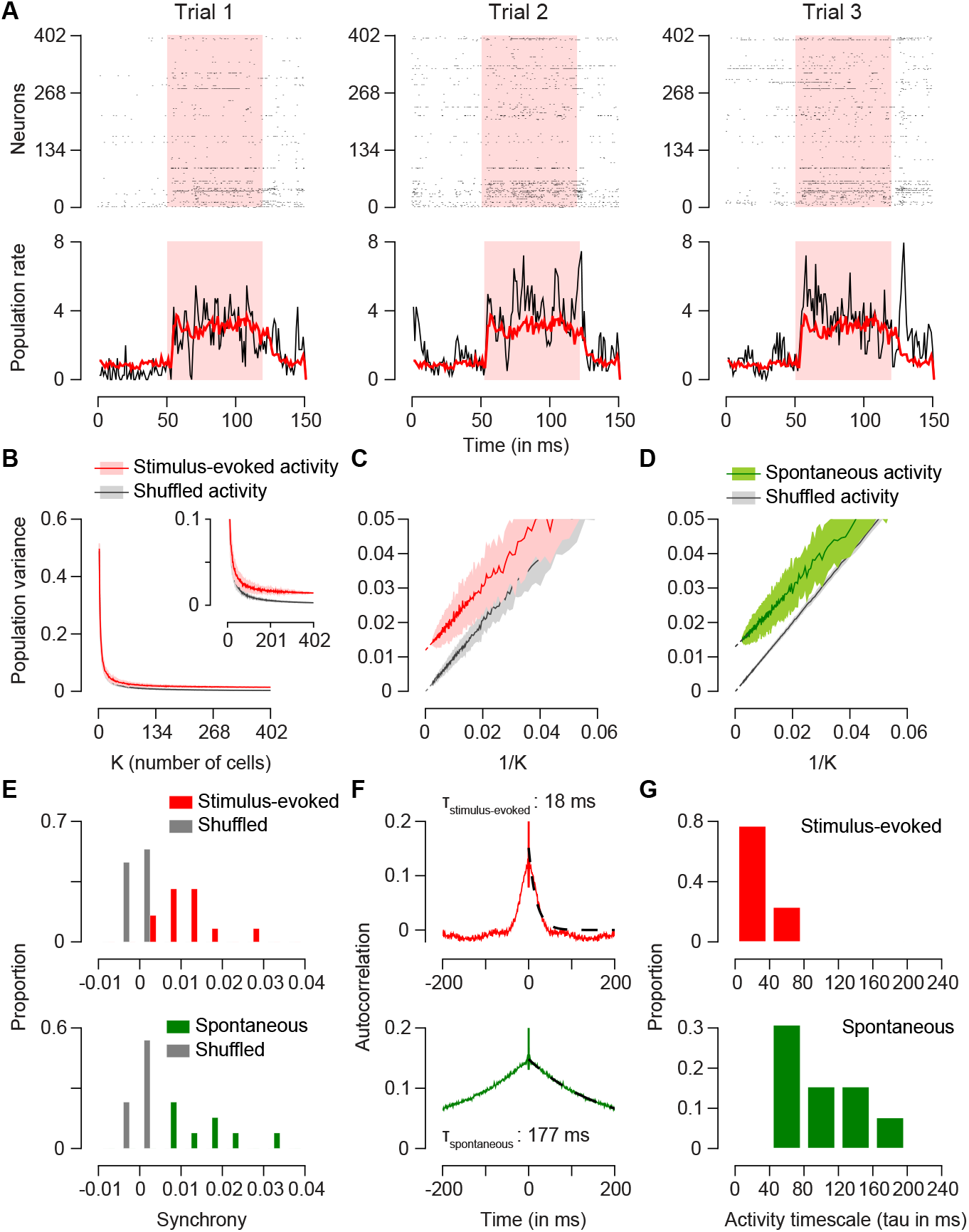
Synchrony in the large-scale extracellular records of cortical networks. A. Raster plots of simultaneously recorded neurons in marmoset primary visual cortex. Each raster shows the response to presentations of the same visual stimulus (shaded region). For each trial the population rate is computed based on the mean response across neurons for a single trial (black traces, bottom) or across all trials (red traces, bottom). B. Left: The population variance (*χ*) is plotted as a function of the number of cells in the population. The standard error (shaded regions) is constructed by randomly resampling the population for a given K (Methods). To generate a null distribution spikes were randomly assigned a trial and *χ* was recomputed (gray). C. *χ* is plotted relative to 1/K. A regression line was fit to estimate the synchrony in the population (43). D. As in C, for spontaneous data. E. The distribution of synchrony estimated from the population variance measure across extracellular records for stimulus-evoked conditions (top, mean = 0.014 ± 0.014) and spontaneous conditions (bottom (mean = 0.027 ±0.025). F. The autocorrelation of the residual population rate during visual stimulation (top) or during spontaneous activity (bottom). Exponentials were fit to the decay of the autocorrelations (dashed lines). G. The distribution of exponential time constant for stimulus-evoked population responses (top) or spontaneous population responses (bottom) (stimulus evoked mean =28.3 ms ± 13.7, spontaneous mean = 126.8 ms +- 102.6).

### *In vivo* synchrony exhibit characteristic time scales

When computed for short time bins (1ms), the *χ*-metric yielded very small spiking correlations, indicating that population responses form of synchrony that is not instantaneous. Rather, populations of neurons exhibit fluctuating responses rates over larger time scales (22, 40, 45-50). We estimated the time scale of these fluctuations by performing an autocorrelation analysis of the population spiking measurements on a trial-by-trial basis and by measuring the time constant of the autocorrelation decay (Fig. 3F). During visual stimulation, the time constant of these fluctuations is broadly distributed with a median value of 29.3 ms (Fig. 3G, mean *τ*_driven_=28.3 ± 13.7). We noticed however, that an exponential decay can be a poor fit to the autocorrelation of population responses. We therefore employed two additional methods to capture the fluctuations time scale. First, we measured the width of the autocorrelation at half-height, which had a mean value of 42 ms ± 16 (51). Second, as using short time intervals to measure these time constants can induce systematic biases, we used a Bayesian method to estimate the fluctuations timescales (52)(Sup Fig. 5). All of these measures indicate that visually evoked fluctuations occur on a time scale between 25 and 50 ms. In contrast, the time scale of spontaneous fluctuations was substantially slower, (Fig. 3F,G bottom, mean *τ*_spontaneous_ = 126.8 ms ± 102.6, width at half height = 241 ms ± 144). Despite these changes in the time scale of population synchrony, we did not find a change in the amplitude of population synchrony, as measured by spiking correlations, between spontaneous and visually-evoked conditions (Fig. 3E, Sup Fig. 3D).

Both the measurements of synchrony amplitude (*ρ*) and time scale (*τ*) were similar in marmoset V1 and area MT. Further, records from mouse visual cortex from datasets collected at the Allen Institute also exhibit similar dynamics, though the amplitude of the synchrony measured in mouse V1 tends to be higher than marmoset V1 (Sup. Fig. 3). In sum, synchrony amplitude, its time scale, and the dependence of time scale on visual drive are common across species in the visual system.

### Weakly synchronous excitatory inputs can generate Poisson-like spiking

We have demonstrated that conductance and large-scale population measurements indicate the presence of synchrony in the network on timescales between 25 and 200 ms. Can such input synchrony also lead to physiological output spiking variability? To answer this question, we generated synthetic conductance traces arising from presynaptic activity with a prescribed degree of spiking correlation. We injected these conductances into neurons *in vitro* (Fig. 4A) and examined the variability of the spiking activity they elicited for the same input rate and input correlation statistics. We found that the spike-count Fano factor was reliably near 1 (Fig. 4C, dark green points, mean Fano factor = 1.2 ± 0.4 s.d., n = 12), across our sample population, consistent with Poisson-like firing observed *in vivo* (Fig. 4D). Moreover, as observed *in vivo (22)*, the Fano factor did not depend on the input firing rate (Fig. 4E) or the bin size used to measure spike counts (Fig. 4F). Irregularity in spiking pattern was also evident in the ISI distributions and CV near 1 (Supp. Fig. 6).

**Figure 4:**
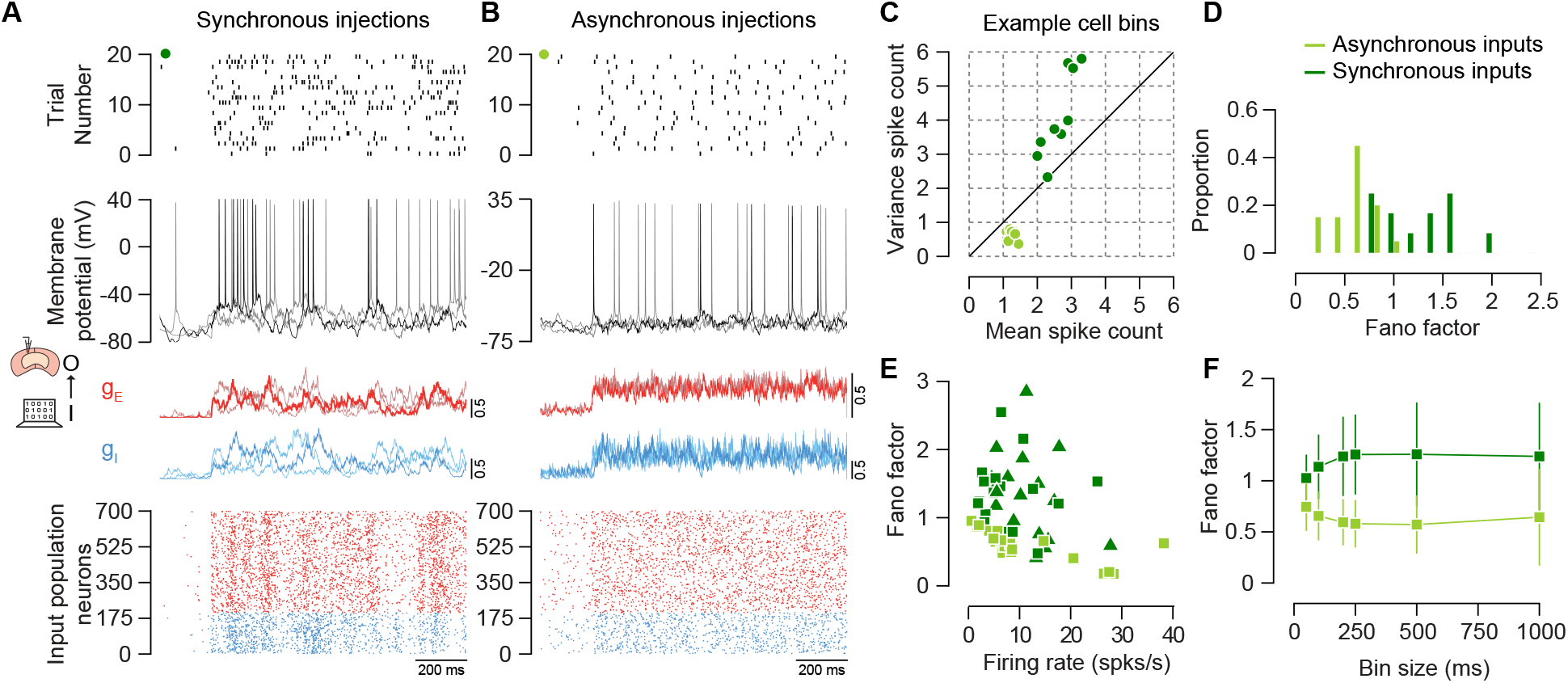
Poisson spiking response emerge from weakly correlated inputs. A. Bottom plot shows a raster of model presynaptic excitatory (red) and inhibitory (blue) neurons generated using a pairwise correlation of 0.030 at a timescale of 50 ms. Those spike trains are used to generate single trials of excitatory and inhibitory conductances (red and blue traces). Three example traces are shown, the light and dark colors are to distinguish the traces. The top two plots show the spike raster and the membrane potential responses of a neuron recorded *in vitro* using dynamic clamp injection of the generated conductances. The first trial is shown in the black trace. B. Same as A, but for conductances generated from an asynchronous network. C. The mean and variance of spike count per bin for the in vitro neuron’s responses in the example trial in black shown in A and B. A sliding 200 ms bin size is used. Dark green points are for the example cell in A with synchronous input and light green points are for example cell in B with asynchronous inputs. D. Distributions of the Fano factor from our set of *in vitro* recordings indicate that responses are slightly above 1 (1.2+/-0.4, n = 12) for synchronous inputs shown in dark green and below 1 (0.6 +/- 0.2 s.d., n = 10) for asynchronous inputs shown in light green. E. The relationship between Fano factor and mean firing rate was not significant for synchronous input (r2 = 0.03, p = 0.3, for asynchronous input R2 =0.5, p = 0.0003). Two different simulated inputs were used for synchronous injections in dark green, one with pairwise correlation of 0.03 indicated by triangles, and another with pairwise correlation of 0.015 indicated by squares. F. The bin size used to count spikes only weakly altered the Fano factor above 100 ms.

To determine whether input synchrony was necessary to generate these variable responses, we also measured the responses of neurons *in vitro* in the absence of input synchrony (Fig. 4B-F). As in the presence of input synchrony, the patterns of action potentials were variable in the absence of input synchrony (Fig. 4B). However, the spike count variance was considerably less than the mean in the absence of input synchrony, resulting in a reduced Fano factor (Mean = 0.6 ± 0.2 s.d., n = 10, Fig. 3C-D). Further, and in contrast to *in vivo* observations (53), the resulting Fano factor was negatively related to the elicited firing rate (Fig. 4E) and to the bin size used for the spike count measurement (Fig. 4F). Low variability also resulted in more regular spiking and smaller CV values for cells injected with asynchronous input. (Supp. Fig. 6)

### Synchrony time scales determine spiking variability *in vitro*

To examine how synchrony timescales impact spiking variability, we consider a statistical model for input synchrony that uses two parameters: a parameter *ρ* that captures spiking correlations at a typical bin size and another parameter *τ* that captures the time constant of the correlation decay. Concretely, we obtained this statistical model by considering that synchronous inputs arise from a fluctuating population-level firing rate, which we model as a nonnegative diffusion process with prescribed autocorrelation time (see Methods). This model generates similar statistics as jittering spikes over a given time window, but allow for an analytical treatment of the amplitudes and time scales of synchrony (28, 46).

We used the above statistical model to generate synaptic inputs with altered synchrony time scale and injected the resulting synthetic conductances into neurons *in vitro* (Fig. 5A). As demonstrated previously, asynchronous inputs generate variable spiking, but with a Fano factor near 0.6. Including synchronous drive at instantaneous time scales (1 ms) increased conductance variability relative to asynchronous drive, but the resulting Fano factor remained low (FF=0.7, Fig. 5A, *τ* = 1ms). The lack of impact of instantaneous synchrony on spiking variability is due the low-pass filtering properties of neurons, which average out fluctuations that are faster than membrane time constant (*τ* ≃ 25ms). Slower input fluctuations pass through the membrane filter, driving larger membrane potential variability for comparable mean level of activity. In keeping with this intuition, we found that longer synchrony time scales leads to systematic increases in the Fano factor of the spiking response of *in-vitro* neurons (Fig. 5A,B). Similar results were obtained using numerical simulations using common electrophysiological models of cortical neurons (54, 55).Injecting synchronous drives revealed a consistent impact of synchrony timescale on spiking variability (Fig. 5C,D). Moreover, we found that recorded and model neurons exhibit similar behavior to changes in input rate, synchrony time scale, and the bin size dependence of the Fano factor as those recorded our experiments *in vitro* (Sup Fig 7,8).

**Figure 5:**
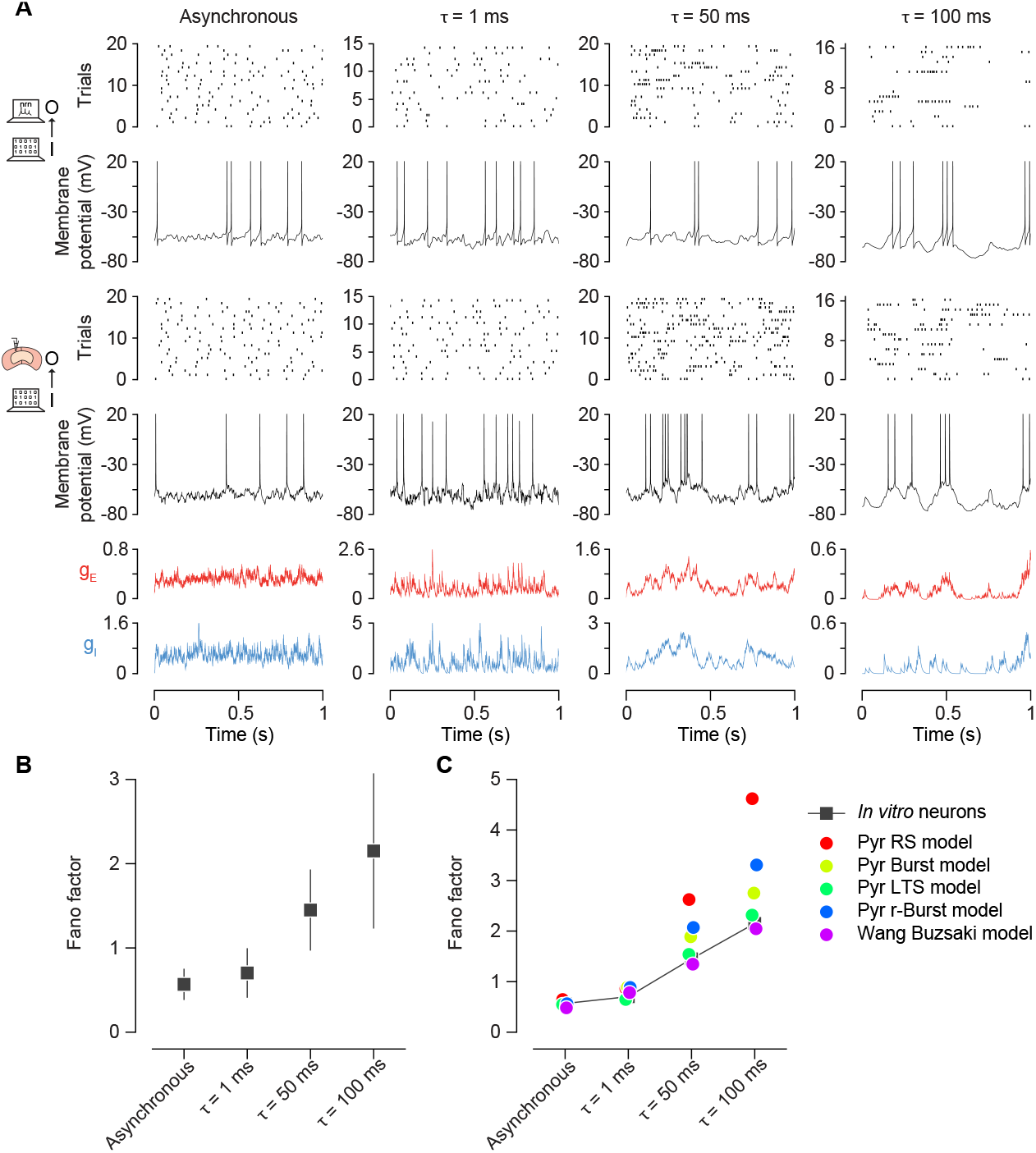
Synchrony timescales alter spiking statistics. A. Spiking activity and membrane potential for model and in vitro neurons for conductances from networks with varying timescale of synchrony are shown. Example conductance traces are shown in the bottom row, in red for excitatory and blue for inhibitory. The middle rows show membrane potential and raster plot upon injecting such conductances in vitro. The top rows show membrane potential and raster plot upon injecting same conductances in model neurons. From left to right, first column shows responses for asynchronous population input, population synchrony with a correlation of 0.015 at 1 ms (second column), 50 ms (third column) and 100 ms (fourth column). B. Measurements of Fano factor for the four conditions presented in A in all in vitro recorded cells. C. As in B, for five different neuron models, indicated by color.

While cortical spiking activity is generally characterized as Poisson, because the Fano factor is near 1, it can also exhibit super-Poisson variability, whereby spike-count Fano factors substantially exceed unit value. Such super-Poisson spiking statistic are found across the cortex during spontaneous activity, whereas input drives tend to quench variability, leading to Poisson-like spiking (4). To determine if we can recover both variability regimes *in vitro*, we injected *in-vivo* conductances recorded during spontaneous (pre-stimulus period) and evoked activity (visual stimulation period). As expected, we found that spiking statistics shifted from super-Poisson during spontaneous drive to Poisson activity during stimulus-evoked drive (Fig. 6A). Our population recording revealed that a key distinction between spontaneous and stimulus-evoked regimes is that the input drives shorten the time scale of synchrony. We hypothesized that this change in synchrony time scale is responsible for the shift from super-Poisson and Poisson spiking. To validate this hypothesis *in vitro*, we injected synthetic conductances for which the synchrony time scale changed from 100 to 50 ms, mimicking a pre-stimulus period followed by a stimulus-driven period (Fig. 6B). To reproduce physiological conditions, we jointly increase the mean input rate and the synchrony timescale when switching from the simulated spontaneous period to the stimulus-driven period. We note, however, that the increase in input drive is not the determinant factor as changing synchrony time-scale alone is sufficient to quench variability (Sup. Fig. 9).

**Figure 6:**
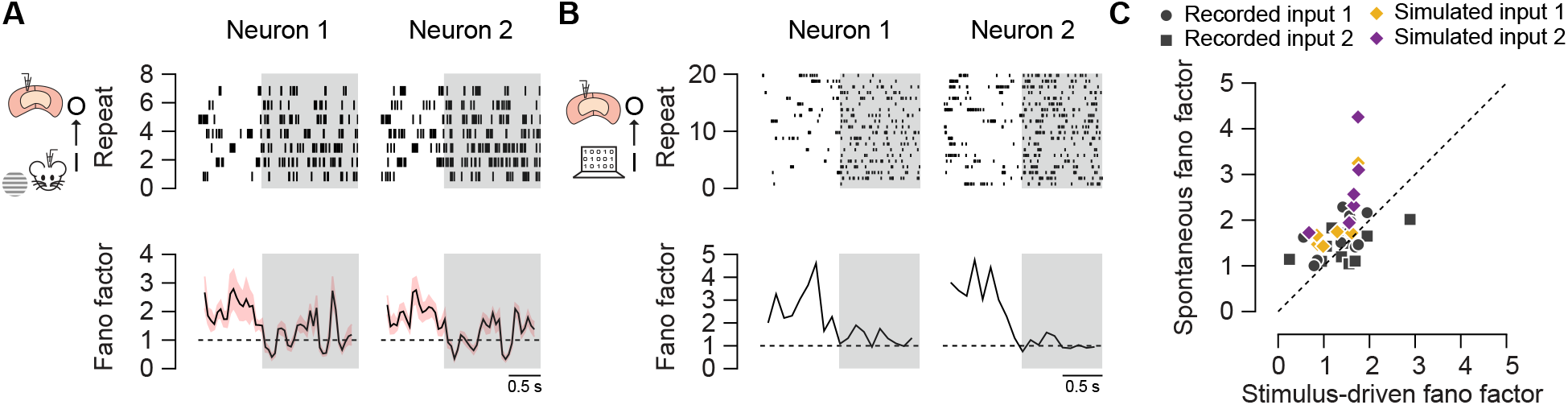
The dynamics of spiking variability. A. Dynamic clamp *in vitro* injections of *in vivo* recorded conductances result in systematic shifts in the Fano factor between the pre-stimulus period (0-750 ms) and the visually-driven period (shaded region, 750-2500 ms). Two example neurons are shown. B. Dynamic clamp *in vitro* injections of modeled population input in which the timescale of synchrony shifts from 100 to 50 ms. C. A comparison of the Fano factor during the spontaneous period and the visually-driven period. Distinct in vivo conductance are indicated by shape (circle, example 1; square, example 2). Distinct modeled synchrony amplitudes are indicated by color yellow (*ρ* = 0.015) and purple (*ρ* = 0.03) diamonds.

### Spiking responses are quasi-deterministic but neuron-specific

Our results show that achieving realistic spiking variability requires synaptic inputs with specific characteristics in terms of synchrony amplitude and time scales. Indeed, we have demonstrated that neurons reliably generate the same spiking patterns when driven by the same conductances (Fig. 1A,B,Fig. 7A,B). At the same time, we find that spiking patterns in response to the same conductances vary significantly from cell-to-cell (Fig. 7A). This suggests that each neuron has its own set of intrinsic properties that sculpt responses. These diverse spiking patterns are especially noteworthy as we neglected the differentiating impact of dendritic integration by presenting conductances to the soma. To quantify the spiking variability due to the heterogeneity of neuronal intrinsic properties, we compared the reliability and precision of single-neuron responses with the responses of distinct neurons driven by the same input (*12*). For repeated injections of same conductance into the same cell, the reliability of spiking patterns across repeats was high (mean = 0.92 ± 0.05, n = 10) and spike times within an event across repeats also showed high precision (mean = 3.9 ms ± 1.3, across = 10, Fig. 7C). We compared these metrics with across-cell measurements by injecting the same conductance in different cells and treating spiking patterns of different cells as a repeat. In this case, the reliability of spiking was considerably lower (mean = 0.72 ± 0.06, across 10 cells) and spike times across cells within an event also showed low precision (mean = 4.6 ms ± 0.6, across 10 cells). These distinct spiking patterns demonstrate how intrinsic properties sculpt responses of neurons differentially. Such response diversity has the impact of reducing the overall spiking synchrony in the population which weakens the spiking correlation of neurons receiving common drive (56).

**Figure 7:**
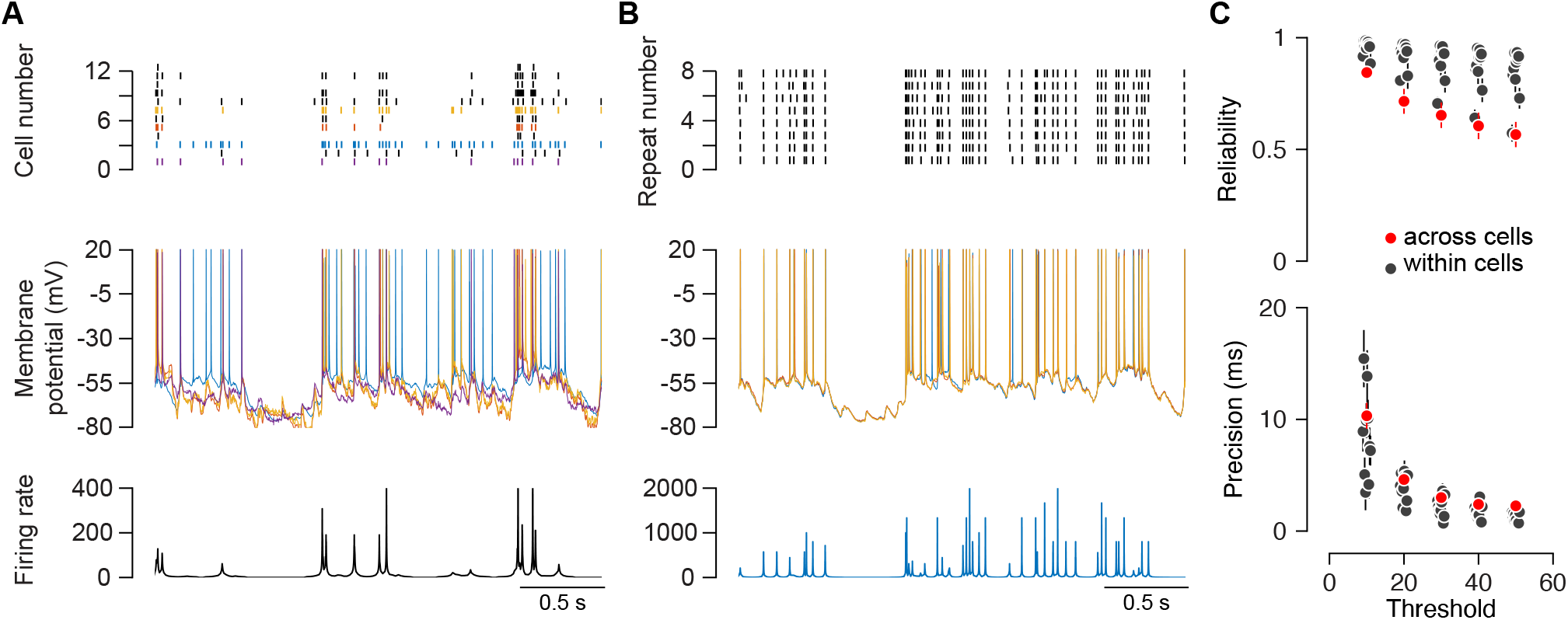
Intrinsic process shape neuronal responses. A. The responses of many neurons recorded *in vitro* to the same conductance input. The population raster shows the spikes of each cell, while the overall firing rate is computed as in Mainen and Sejnowski (1995). Membrane potential traces are shown for four of the neurons in the sample population. B. The responses of a single neuron (4) to the same stimulus presented repeatedly. Firing rate and membrane potential as in A, but for this single neuron. C. The reliability of population response (red) is always lower than individual neurons (black) and declines steadily with the threshold rate. D. Individual neurons are more precise than the population.

## Discussion

We have demonstrated that irregular spiking observed in single cells *in vivo* results from external synaptic drive. Our experimental and modeling results revealed that synchrony is an essential determinant of cortical variability, which poses several challenges to existing theories.

To support this claim, we demonstrated a network-level origin for cortical variability via two key observations at the single-cell level. First, we showed that neurons respond reliably to repetition of the same physiological synaptic inputs that were recorded *in vivo*, as suggested by previous *in-vitro* studies that used synthetic input (12). Second, we found that spiking variability is fully accounted for by the fluctuations of the synaptic drive that a neuron receives in response to the same sensory stimulus. This supports the idea that neurons faithfully respond to variable inputs, whose fluctuations originate from network properties rather than intrinsic sources. Further, we have established that cross-trial fluctuations in the excitatory drive are sufficient to evoke variable spiking responses. This suggests that, at the single-cell level, balancing strong excitation and inhibition in the inputs is not a requirement for spiking variability to emerge.

We argue that the physiological synchrony in networks generates input fluctuations that result in irregular spiking. To support such a role for synchrony, we followed four lines of evidence. First, statistical analyses of conductance traces recorded *in vivo* shows that the strength of their fluctuations can only be explained by some degree of synaptic input synchrony. Second, using a population correlation analysis of large-scale spiking recordings we quantified the presence of spiking correlations. Third, we developed models that generate inputs with various degree of synchrony. With these models, we showed that the level of spiking correlation measured in our experiments yields synthetic conductance traces with physiological levels of fluctuations. Fourth, injecting neurons with these synthetic conductances demonstrates that input synchrony is sufficient to drive physiological spiking variability *in vitro*.

Our measurements identify the time scale of synchrony as a key component controlling spiking variability. Shifting synchrony timescales from slow to fast, as occurs at the transition from spontaneous and driven states, reduces the degree of spiking variability from super-Poisson to Poisson, as has been observed across cortical areas. The distinction between super-Poisson and Poisson spiking regime is thus explained by frequency modulation rather than amplitude modulation (4).

It remains unclear how such synchrony emerges within cortical networks. We presented evidence that synchrony not only exists at the level of the cortex, but in the afferent inputs from the thalamus (Sup. Fig 3), suggesting that cortical networks may only need to maintain and modulate synchrony. Correlated activity varies with internal state (e.g. attention), suggesting that there are multiple stable synchronous network states. Network models composed of multiple clusters of interconnected neurons, as observed *in vivo* (57-61), can generate metastable dynamics with slow synchronous network fluctuations, whose timescales are modulated by input drives (62). Mechanistically explaining the origins of cortical variability likely hinges on understanding the stable emergence and maintenance of these synchronous fluctuations in structured spiking networks (6, 62-64). This will probably require considering network models beyond classical approximations that neglect non-Gaussian, correlation-based, correction terms (65-67).

## Methods

All marmoset and mice experiments were conducted with the approval of The University of Texas at Austin and University of Nevada at Las Vegas Institutional Animal Care and Use Committees.

*In vivo* physiology procedures:3 male and 1 female marmoset was used in the current study. These animals had chambers implanted over primary visual cortex or area MT. Surgical procedures were similar to previous descriptions (68). Custom-made headpost and chambers were affixed to the skull in a sterile anaesthetized procedure. Throughout the procedure, the body temperature was maintained at 36-37°C and the heart rate, SPO2 and CO2 were monitored. Animals were placed in stereotaxic frames, circular craniotomies were performed on the intended chamber location identified using stereotaxic coordinates, chambers and the headpost were placed and the dura was removed. An implant from dental acrylic was built around the headpost and chambers, covering the remaining exposed skull. The skin around the implant was affixed to the implant using Vetbond. The animals were then returned to the cages after recovery from anaesthesia.

Chamber design: The chamber consisted of 4 parts. The outermost part of the chamber was a ring of height 1.6 mm and of diameters ranging from 5-7mm. This ring had 1 mm long thin feet that were inserted inside the skull following craniotomy. The second piece was a thin chamber nut (thickness 1.5 mm) that was screwed on the outside of the chamber ring and rested on top of the skull. This assembly was further sealed using Metabond (Parkell, New York). A removable imaging well was screwed on the inside of the chamber ring. The well consisted of a metal insert to which a coverglass was attached at the bottom. A thin cap (1 mm) was screwed on top of the chamber ring to close it.

Behavioral training and experimental control: After recovery from surgery, marmosets were habituated to head fixation and trained to fixate visual targets. Experimental control was provided by the Maestro software suite, which collected eye movement data, controlled visual stimulation, and provided juice reward (https://sites.google.com/a/srscicomp.com/maestro/).

Population recordings: Large scale population recordings from V1 and area MT data were collected using Neuropixels 1.0 probes. We used an IMEC PXIe acquisition module mounted on a National Instruments (NI) PXIe chassis (PXIe-1071) with NI PXIe-8381 and NI PCIe-8381 for remote control. Voltage signals were recorded at 30 kHz from 384 channels using SpikeGLX. Waveforms were first automatically sorted using Kilosort and then manually curated using the phy software (69). Large-scale population recordings in mice were downloaded from the publicly available dataset at the Allen Institute (70).

*In vitro* physiology procedures: Mice and marmosets underwent cardiac perfusions with ice-cold saline consisting of (in mM): 2.5 KCl, 1.25 NaH2PO4, 25 NaHCO3, 0.5 CaCl2, 7 MgCl2, 7 dextrose, 205 sucrose, 1.3 ascorbicate acid, and 3 sodium pyruvate (bubbled constantly with 95% O2/5% CO2 to maintain pH at ∼7.4). The brain was removed and sliced into 300 µM sections containing V1 region of cortex or temporal association cortex were made using a vibrating tissue slicer (Vibratome 300, Vibratome Inc). The slices were placed in a chamber filled with artificial cerebral spinal fluid (aCSF) consisting of (in mM): 125 NaCl, 2.5 KCl, 1.25 NaH2PO4, 25 NaHCO3, 2 CaCl2, 2 MgCl2, 10 dextrose, 1.3 ascorbic acid and 3 sodium pyruvate (bubbled constantly with 95% O2/5% CO2) for 30 minutes at 35°C and then held at room temperature until time of recording.

### In vitro Electrophysiology

Slices were placed in a submerged, heated (32-34 C°) recording chamber and continually perfused at 1-2 ml/min with aCSF (in mM): 125 NaCl, 3 KCl, 1.25 NaH2PO4, 25 NaHCO3, 2 CaCl2, 1 MgCl2, 10 dextrose, and 3 sodium pyruvate (bubbled constantly with 95% O2/5% CO2). Ionotropic glutamatergic and GABAergic synaptic transmission were blocked with 20 µM DNQX, 25 µM D-AP5, and 2 µM gabazine. Neurons were visualized with a Zeiss AxioExaminer under 60x magnification. All drugs were obtained from Tocris, Abcam pharmaceutical, or Sigma and prepared from a 1000x stock solution in water.

Whole cell recordings were made using a Dagan BVC-700 amplifier and custom written acquisition software using Igor Pro (WaveMetrics) or Axograph X (Axograph). Data were sampled at 20-50 kHz, filtered at 3 kHz, and then digitized by an InstruTECH ITC-18 interface (HEKA). The internal recording solution consisted of (in mM): 135 K-gluconate, 10 HEPES, 7 NaCl, 7 K2-phosphocreatine, 0.3 Na-GTP, 4 Mg-ATP (pH corrected to 7.3 with KOH). Recording electrodes were pulled using Flaming/Brown puller (Model P-97, Sutter Instruments) from borosilicate glass (outer diameter 1.65 mm, World Precision Instruments) and had an open tip resistance of 4-6 MΩ. Series resistance was compensated using the bridge balance circuit and was monitored throughout the experiment. Experiments were discarded in series resistance exceeded 35MΩ.

Dynamic clamp experiments were performed using a Teensy 3.6 microcontroller that converted excitatory and inhibitory conductance commands with the records of membrane potential into current at a high rate (100 kHz) (Desai et al. 2017). Excitatory and inhibitory reversal potentials were set to 0 and -80 mV, respectively. Action potentials were identified by extracting the times at which membrane potential exceeded a threshold voltage.

Conductance commands: Conductances injected *in vitro* were either from measurements made *in vivo* (Tan et al.) or using generated synthetically at a timescale of 8kHz. The gain of the i*n vivo* conductances was adjusted equally during experiments to generate approximately 7 spikes/sec, except for those experiments in which the conductance gain was systematically adjusted (Sup. Fig. 2). The relative gain of excitatory and inhibitory conductances was not varied, apart from those experiments in which inhibitory conductance was set to 0. Baseline holding currents at either the reversal potential for excitation or inhibition were computed from the bottom fifth percentile of the distribution of current values. These baseline currents were subtracted from the current traces and converted into conductance traces, which were used as commands to the dynamic clamp system.

Synthetic conductances were generated as shot-noise traces by exponentially filtering the spiking patterns of 500 excitatory neurons and 200 inhibitory neurons characterized by a rate, correlation, and correlation time scale. We simulated synchronous spiking activity via as a doubly-stochastic procedure, whereby the common neuronal spiking rate is prescribed as a random Cox-Ingersoll-Ross (CIR) process. Such a process is governed by

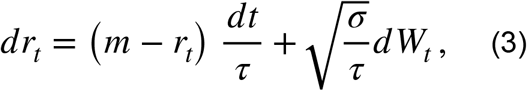

where *m* is the mean firing rate, *τ* is the correlation time, *σ* is the noise coefficient, and *W* is a Brownian motion. Then, to simulate spiking inputs from *K* inputs, we sample the number of active inputs in each time bin of duration Δ*t* by sampling a Poisson random variable with parameter *KrΔ t*, where *r* refers to the (average) value of the fluctuating rate in that time bin. Given *m* and *τ*, we choose *σ* to achieve the level of experimentally measured level of spiking correlations (the larger *σ*, the larger the spiking correlation *ρ*). In our simulations, Δ*t* =1/8000s, while *m, ρ* and *τ* were varied. The spiking correlation *ρ* was varied by choosing *σ* when measured in time bins of 15 ms duration. The degree of correlation between excitatory and inhibitory inputs was also varied by considering that each type of inputs were driven by two separate CIR processes *r*_*e*_ and *r*_*i*_. This varying degree of correlation was obtained by allowing for *r*_*e*_ and *r*_*i*_ to be driven by a shared noise component in equation (3): 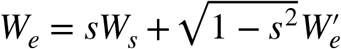 and 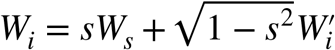 where *s* denotes the fraction of shared noise, *W*_*s*_ denotes a shared Brownian motion drive and where 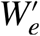 and 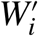 denote independent private Brownian motion drives. Input spikes were passed through exponential decays of either 5 ms (excitatory neurons) or 10 ms (inhibitory neurons). The overall gain of inhibitory drive was set to twice the excitatory gains, which matches the difference in conductance gain measured between excitatory and inhibitory conductances.

Neuron simulations: We used the single compartment models of regular-spiking pyramidal cell, bursting pyramidal cell, repetitive bursting pyramidal cell, and low-threshold spiking (LTS) pyramidal cell from Pospischil et al. 2008 (55) as well as the model of interneuron of Wang-Buszaki (54). We downloaded these five models from the ModelDB repository and used the simulation environment Neuron (71, 72). The parameters were not adjusted. Simulations were either performed at 8 kHz, for the neurons from Pospichil et al., or at 160 kHz for the Wang-Buzsaki neuron. The conductance of the external inputs was generated using the same method for those cells recorded *in vitro*. As for *in vitro* experiments, action potentials were extracted by identify the time at which membrane potential exceeded a voltage threshold.

Population variance: Measurements of *χ* were made from simultaneously recorded populations of neurons. The number of neurons included in the *χ* measurement was systematically varied from K= 2 to the total number of neurons (K_tot_)in the recorded population. For each K, K neurons were selected randomly and the population variance was computed by measuring the variance of the population rate across trials relative to the summed variance of the individual neurons (equation 2).

For each subpopulation size K, K neurons were selected 50 times randomly. Both the mean and 95% confidence intervals of *χ* were computed from mean and standard deviation of this distribution. We then shuffled the spikes randomly between the trials, to disrupt any trial-by-trial covariance, and performed the same analysis (51). To measure the population variance as K approaches infinity was extracted from the y-intercept of the linear regression using the log-transformed *χ* values from the 1/K_tot_ to 1/ (K_tot_*0.5).

Synchrony time course: Estimates of the timescale of population synchrony were made using two different procedures. First, we estimated the synchrony time course from the decay in the autocorrelation of the residual population response. We measured by the population response from the summed activity of neurons in a given trial, and computed the residuals from the difference in the individual trial populations from the population responses averaged across all repeats of the same stimulus. The autocorrelations of these residual responses from each trial and across conditions were averaged and an exponential time constant was fit to the average autocorrelation. The second method we used to estimate synchrony time scale was to employ the method described in Zeraati et al 2022, which uses a generative model based on Ornstein-Uhlenbeck processes.

Measurements of precision and reliability were based on the procedure outlined in Mainen and Sejnowski, 1995 (12). PSTHs were generated either for neuronal populations, by integrating the responses of distinct neurons to the same conductance injection, or by integrating the responses of a single neuron to injection of the same conductance. An adaptive filter was applied to the average responses which was centered on each time bin and widened until half of the responses contained a spike or the bin width reached 100 ms. Events were then identified as by firing rate crossing a threshold level of firing rate, which was varied systematically in steps of 10 from 10 to 50 spikes/s. Reliability is defined as the fraction of spikes that occur within these events, whereas precision is defined as the standard deviation of spike times within an event.

## ACKNOWLEDGMENTS

We thank Carl van Vreeswijk for his thoughtful advice on this project. We thank Carrie Barr, Vy Nguyen and Mason Antin in the Priebe laboratory for lab management and animal care. D.H. would like to thank Carole Levenes for discussions and Yonatan Loewenstein, Edmond and Lily Safra for Brain Research at the Hebrew University of Jerusalem for hospitality.

Funding: D.H. was funded by ANR-14-NEUC-0001-01 ; ANR-17-NEUC-0005, D.B. was funded by NIH R01 MH131317, T.T. was funded by NSF-Career Award: DMS-2239679, CRCNS Award: DMS 2113213, N.J.P was funded by NIH R01 EY025102 and NS120562, R.T.O by a CPS Training Fellowship.

## Author contributions

Conceptualization: JJP, DH, TT, DB and NJP

Experimentation: JJP (marmoset), DB (in vitro), RTO (mouse neuropixels)

Analysis: JJP and NJP

Simulations: JJP, TT and NJP

Analytical derivations: DH and TT

Writing and editing: JJP, DH, TT, DB and NJP

## Supplementary files

1. Supplementary figures: 1-9
2. Analytical derivations

## Supplementary material

**Supp. Fig. 1:**
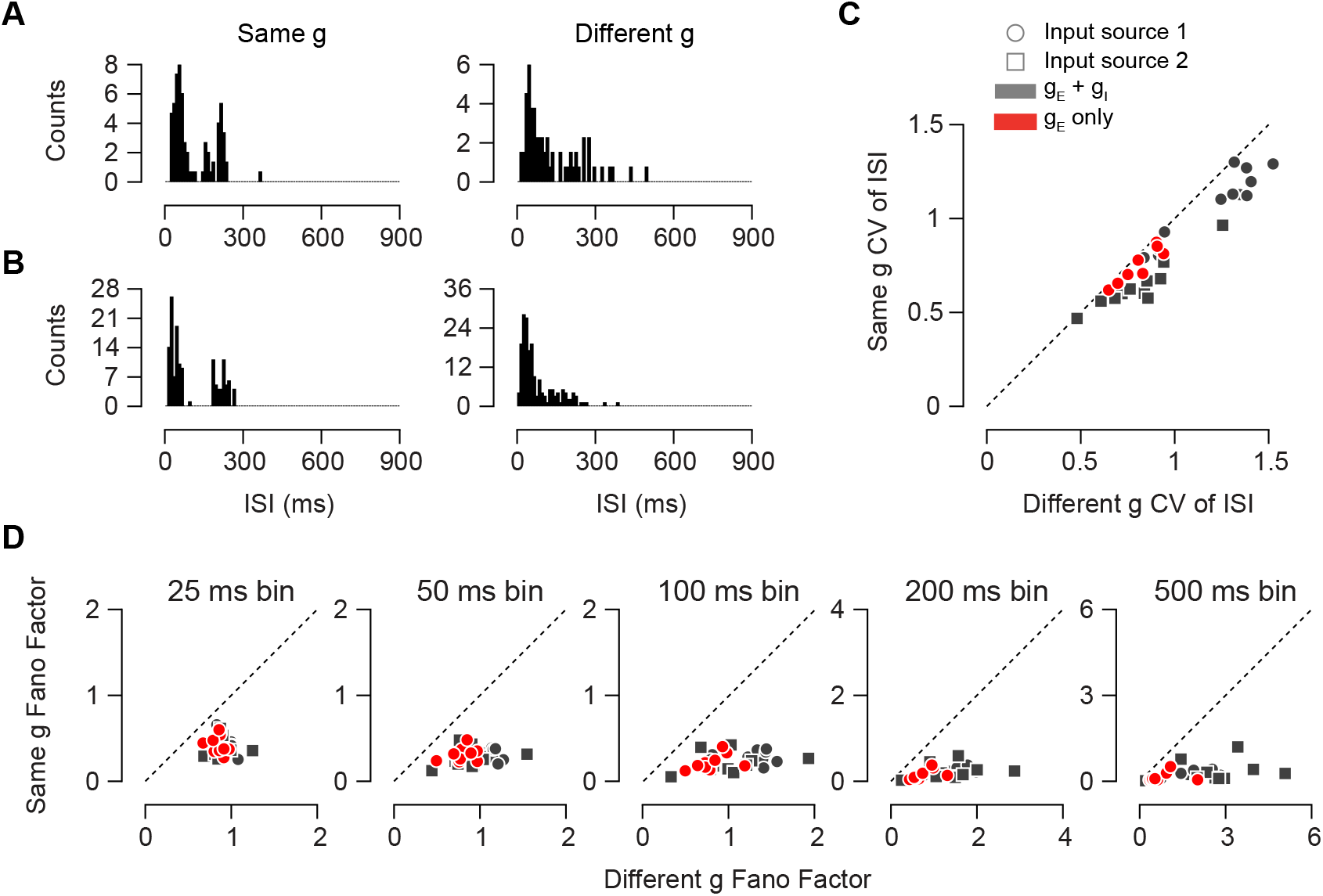
CV and bin size dependence of Fano factors. A and B.Distributions of the inter-spike interval are shown for the example cell in Fig. 1A and C respectively. The cell in A is injected with g_E_ + g_I_, while cell in B is injected with g_E_ alone. Left and right columns are output from same g and different g injections respectively. C A comparison of CV of the ISI distribution for all cells which input conductance is the same or different. D. A comparison of the Fano factor using different bin sizes. Markers are same as in C.

**Supp. Fig. 2:**
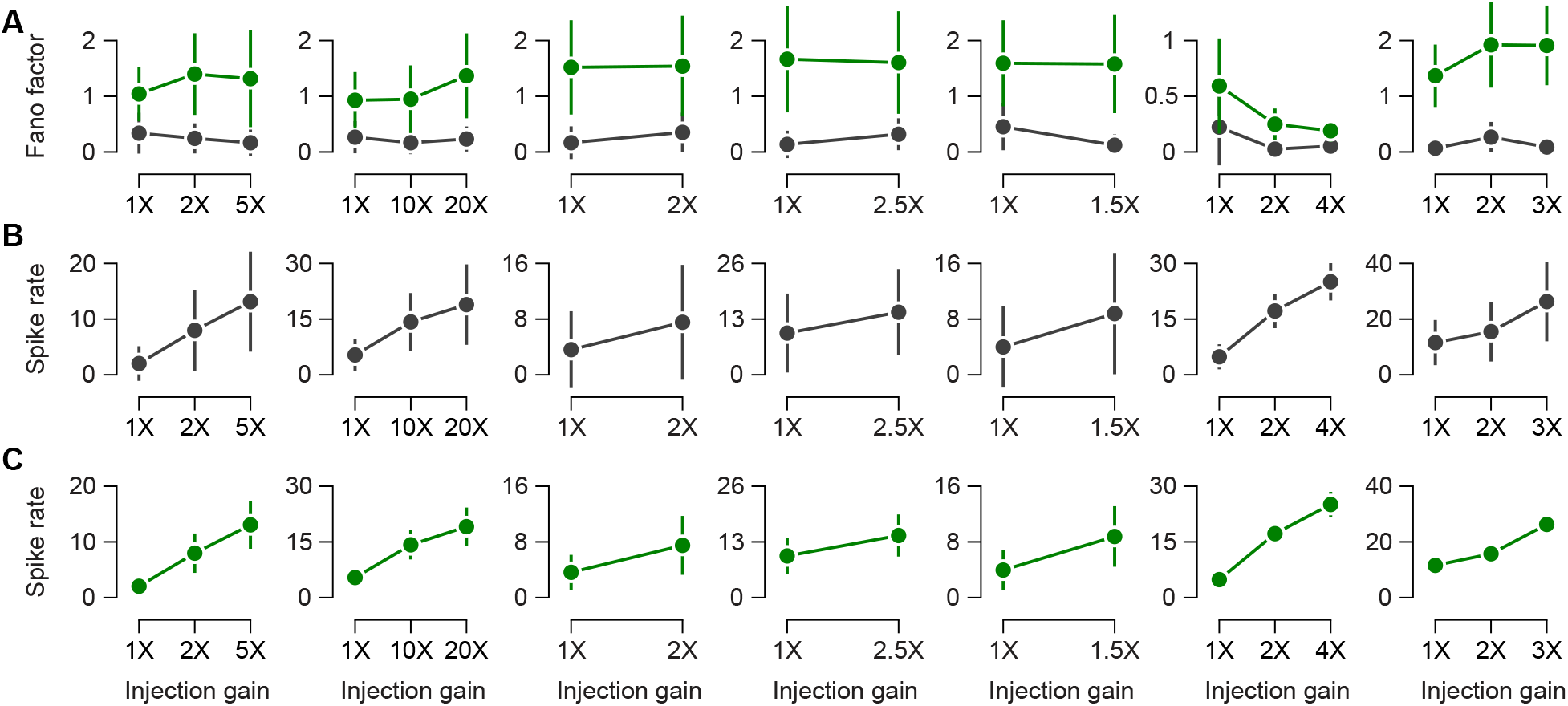
Effect of changes in gain on Fano factor and firing rate. A. The Fano factor for changes in excitatory and inhibitory conductance gains example cells. Each column is a separate cell. Green points are output from different g injections and gray points are output from same g injections. B. The changes in spike rate as a function of gain for the cells in A, for same g injections. C. The changes in spike rate as a function of gain for cells in A, for different g injections.

**Supp. Fig. 3:**
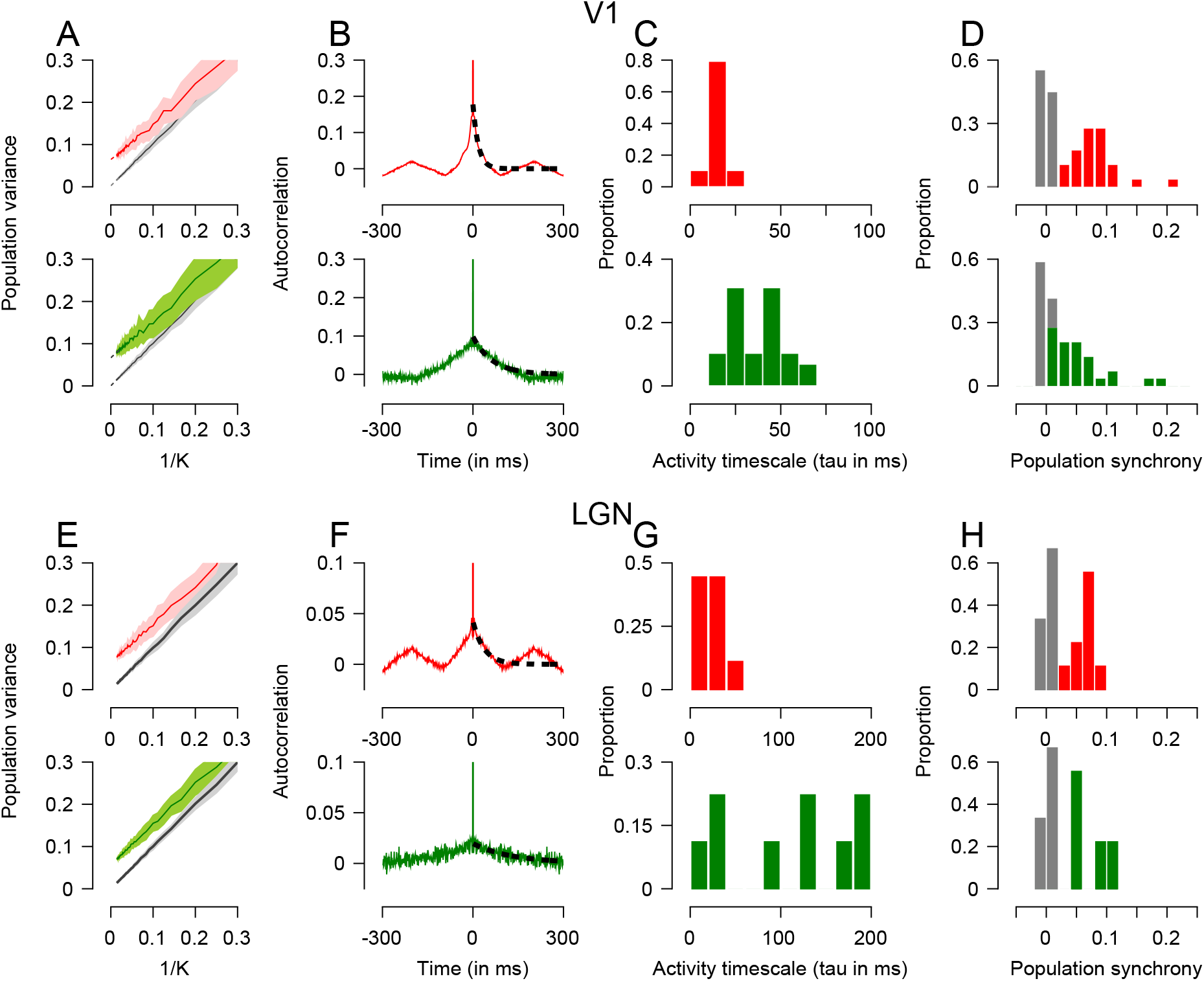
Synchrony in large scale extracellular recordings from mouse cortex and visual thalamus (LGN). A. Shows population variance as a function of number of cells during stimulus-evoked (top) and spontaneous (bottom) conditions for an example session. B. Show autocorrelation of population activity for the session in A. The dotted line shows the fitted exponential decay. C. Distribution of population activity timescale (tau) across all sessions. D. Distribution of synchrony across all sessions. E-H, as for A-B, but for thalamic recordings. Data is from Allen Institute (70).

**Supp. Fig. 4:**
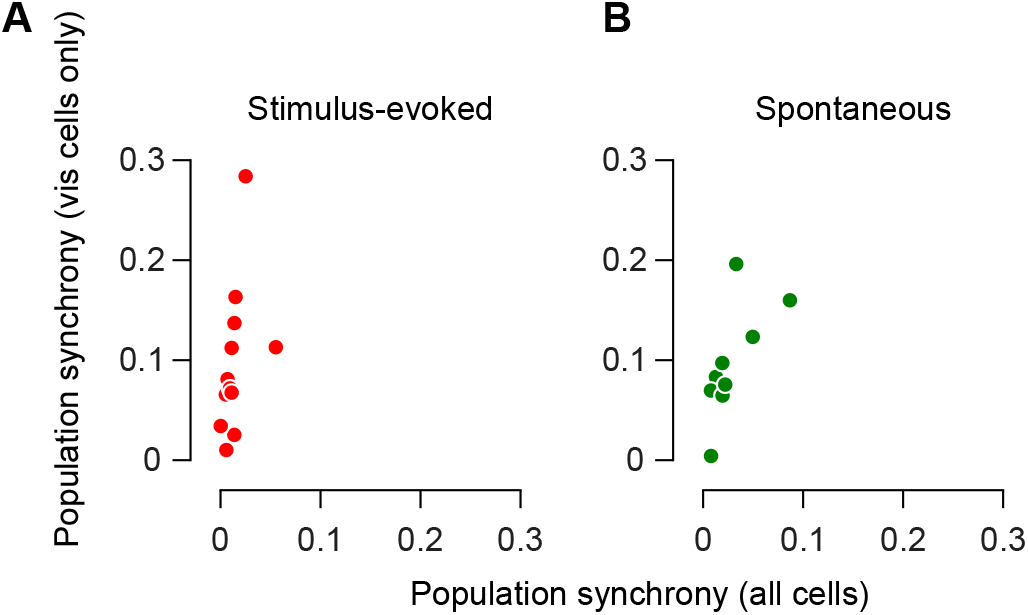
Synchrony comparison for sub-population of visually responsive cells. A and B. Show how population synchrony changes for marmoset cortical populations upon only including the visually responsive cells. A shows data for stimulus-evoked synchrony and B is for spontaneous synchrony.

**Supp. Fig. 5:**
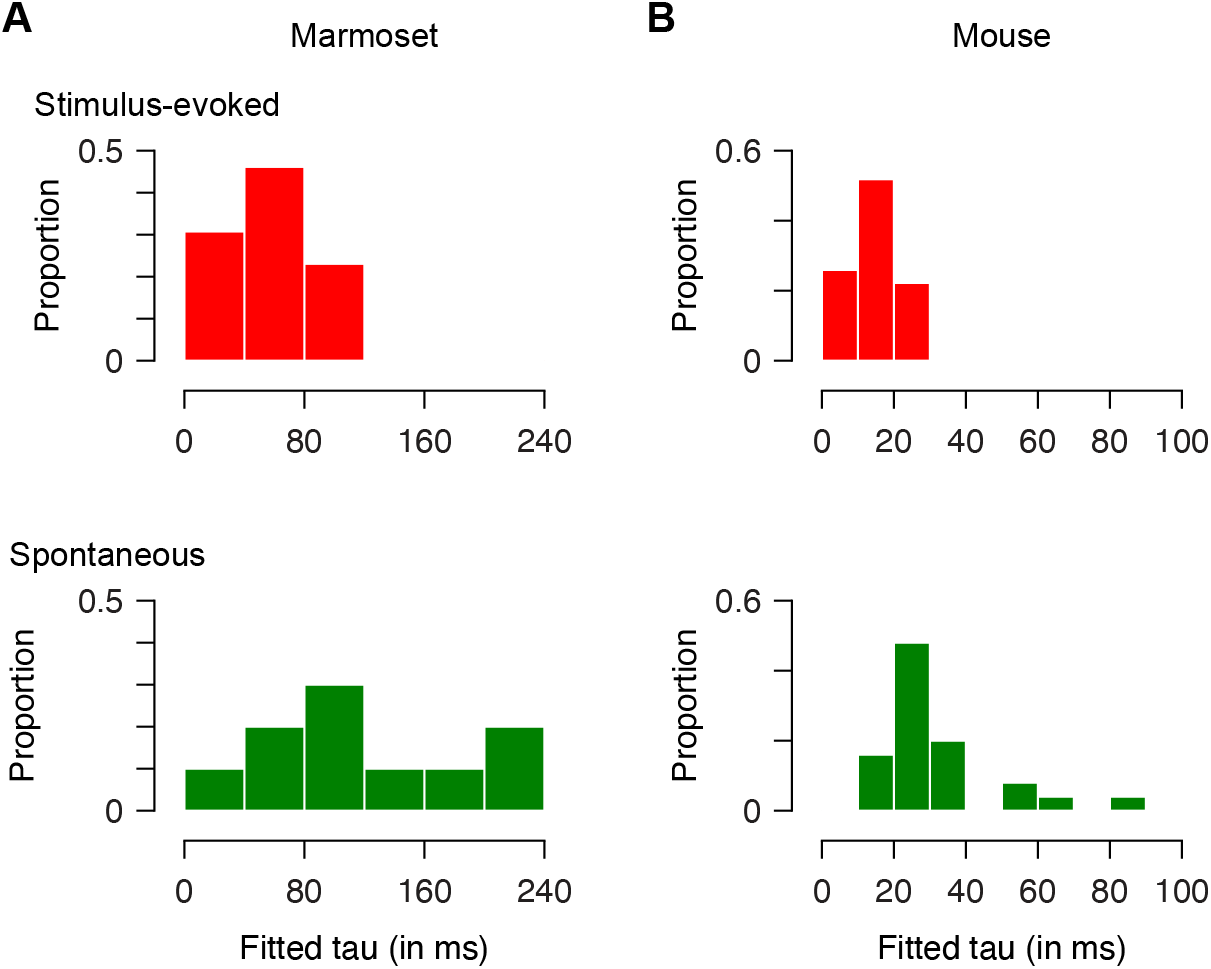
Estimation of activity timescales using abcTau method. A. Shows distribution of timescales (tau) for marmoset cortical populations using the abcTau method. Top row is stimulus evoked timescale, bottom row is spontaneous timescale. B. Same as A, for mouse cortical populations.

**Supp. Fig. 6:**
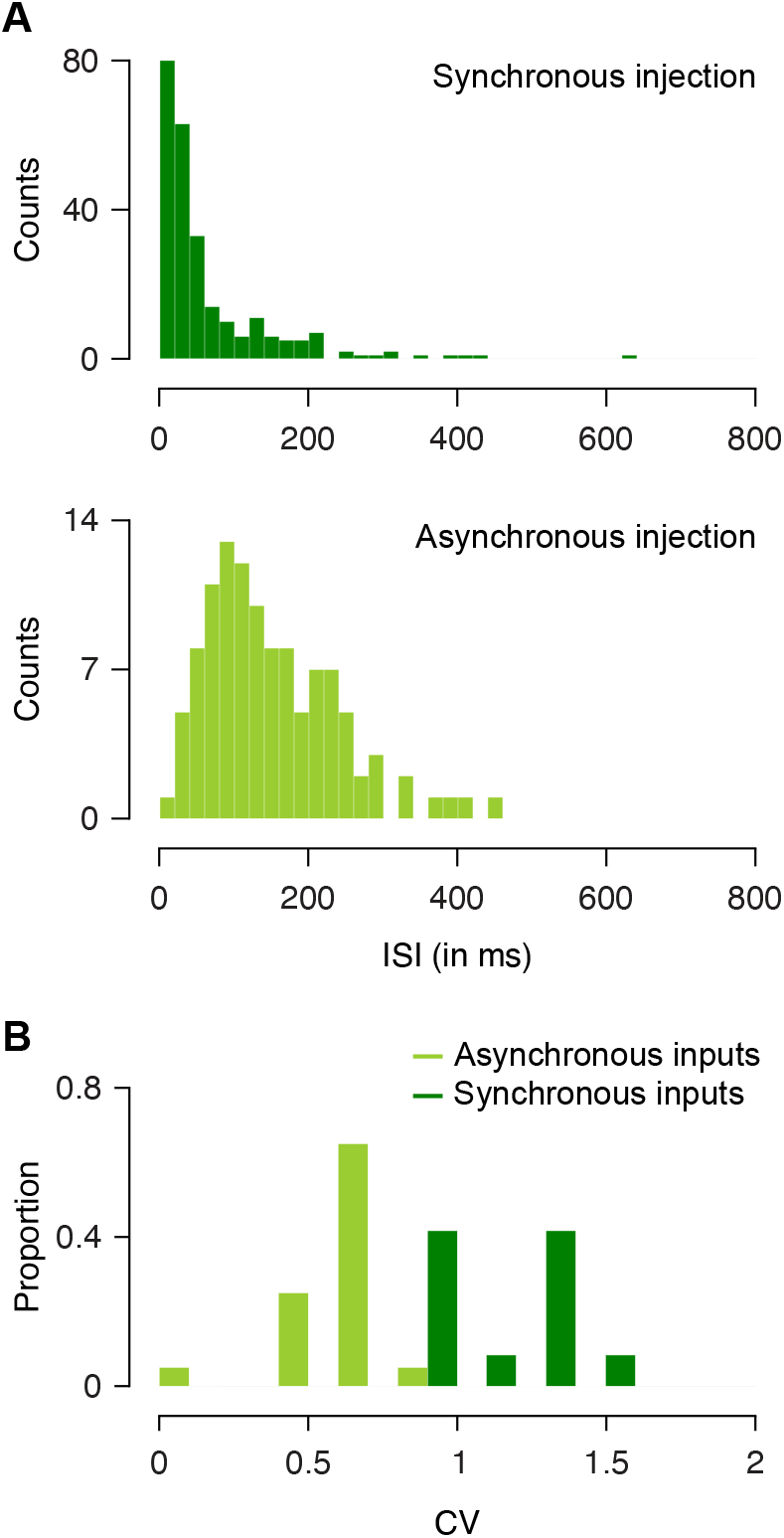
ISI distribution and CV for synchronous vs asynchronous input. A. Shows ISI distributions for synchronous (top) and asynchronous (bottom) injections in example cells shown in Fig. 4 A and B. B. Distribution of CV across all cells recorded with synchronous (dark green) and asynchronous (light green) inputs.

**Supp. Fig. 7:**
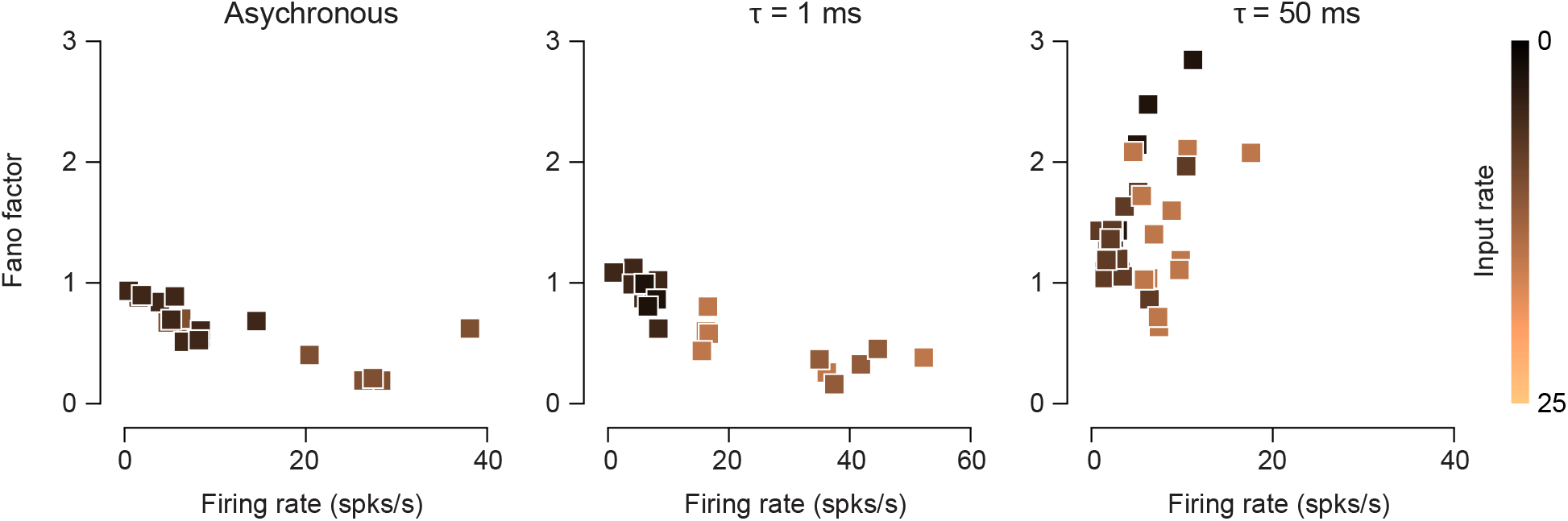
Dependence of the Fano factor on output firing rate and input rate under varying input correlation timescale conditions. Each point is the Fano factor of a cell injected with input with synchrony at specified time-scale (panel titles) and is color coded by the input rate used for that point. Same cells were injected with multiple input rates.

**Supp. Fig. 8:**
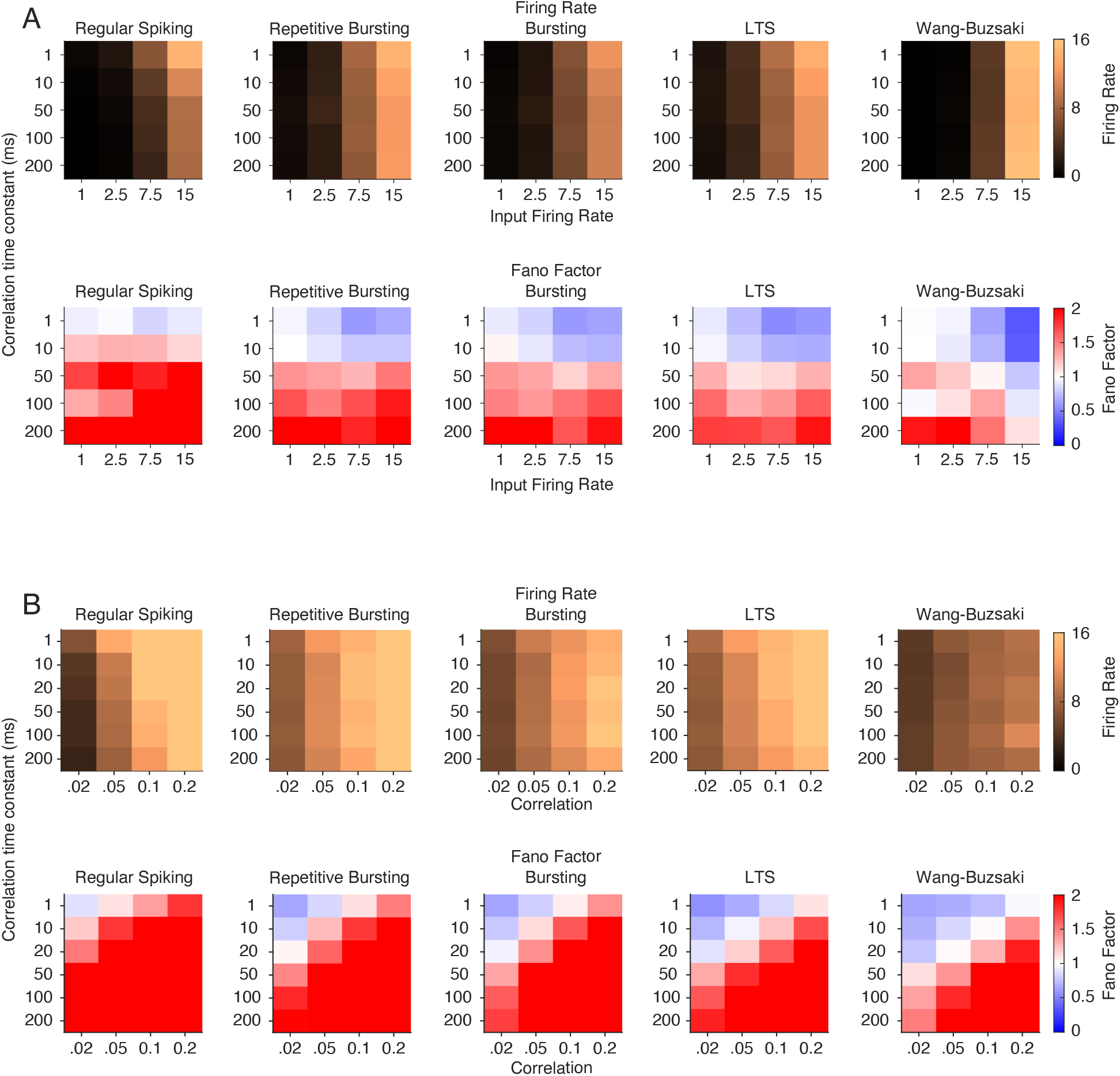
Variations in Fano factor and output firing rate with synchrony timescale, input rate, and correlation in simulations. A. The firing rate (top) and fano factor (bottom) are shown for the 5 types of model neurons. The synchrony time constant and input firing rate were systematically altered for a pair-wise correlation of 0.02. B. As in A, but the synchrony time constant and pair-wise correlation were systematically varied. The input firing rate was set to 7.5 Hz.

**Supp. Fig. 9:**
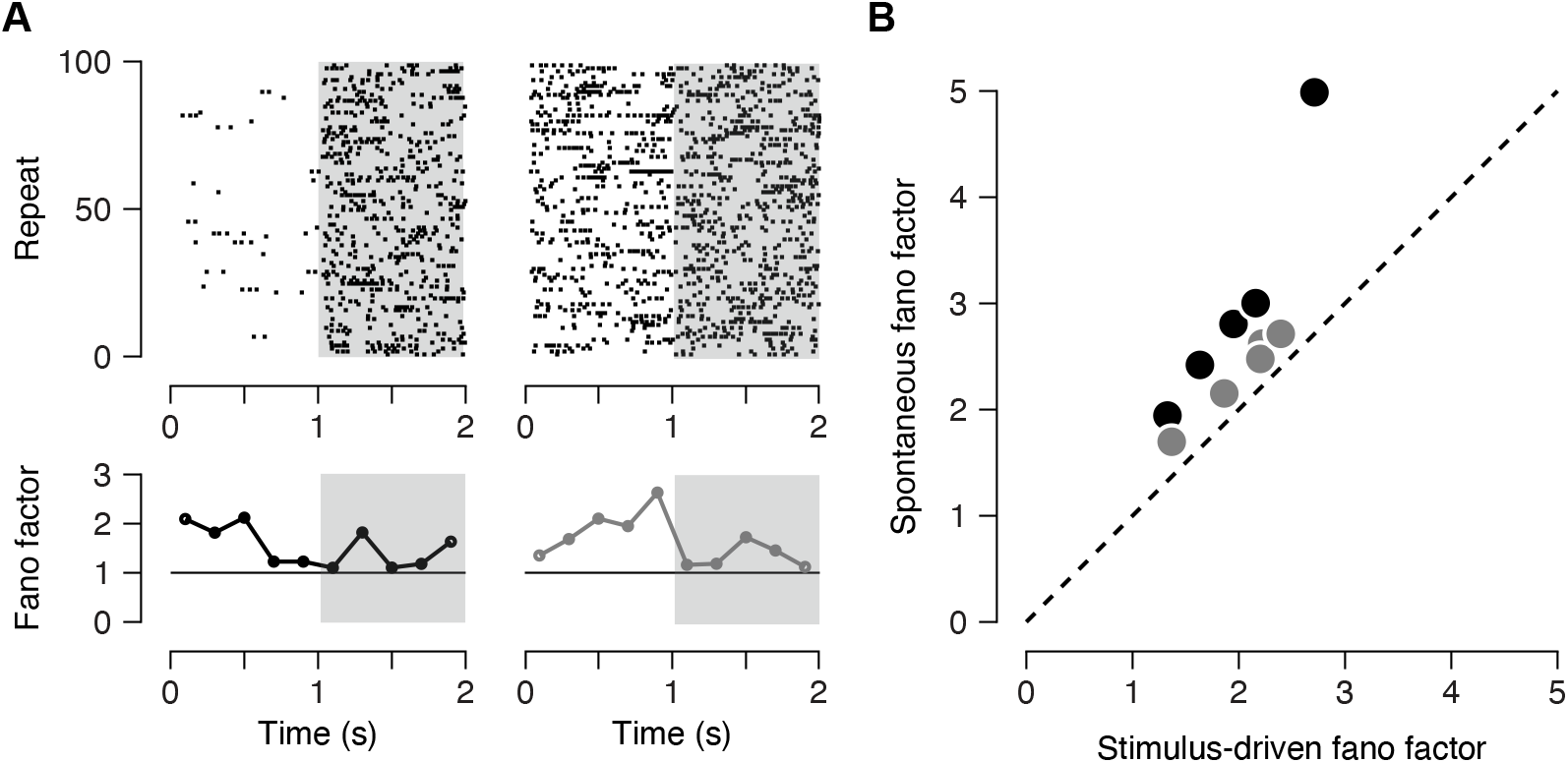
Variability quenching in simulated neurons. A. Top row shows raster for an example simulated neuron. Gray shaded region is when time-scale of simulated input shifts from 100 ms to 50 ms (same as in Fig. 6B). Bottom row shows the Fano factor of the model cell in 200 ms bins. The left column is when firing rate is allowed to change from simulated spontaneous (1 spk/s) to stimulus evoked (7.5 spks/s) period. Right column is when firing rate is held constant through the 2 s trial (7.5 spks/s). B) Shows spontaneous and stimulus evoked fano factor comparisons for 5 types of neuron simulation models. Black points are for input with firing rate changes, gray points are for constant firing rate.B. The mean Fano factor for each model cell in the driven (synchrony time scale = 50 ms) and spontaneous periods (synchrony time scale = 100 ms) for each model cell. Changes in Fano factor were measured when the input firing changed from 1 to 7.5 spks/s (gray symbols) or when the input firing rate was fixed (black symbols).

## Supplementary information

### 1 *χ*-metric estimates for spiking correlations

In this section, the scaling behavior the *χ*-metric with respect to the number of considered neurons *K*. To do so, we consider a series of spiking statistical models under increasingly realistic assumptions.

#### 1.1 Temporally independent, homogeneous case

##### 1.1.1 Bernoulli counts

Consider some spike train data *X*_*i,m*_, 1 ≤ *i* ≤ *K*, where *i* is the neuron index and *m* the time index. Assuming stationarity, we can neglect time dependence and disregard *t*. We also assume exchangeability and allow for nontrivial correlation. Let us define

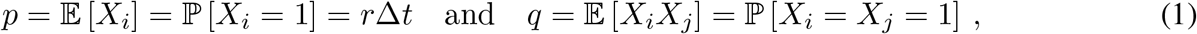

where *p* is the probability to find a spike in a bin, where *r* denotes the spiking rate, and where *q* = *p*^2^ in the absence of correlations. The spike correlation is defined as

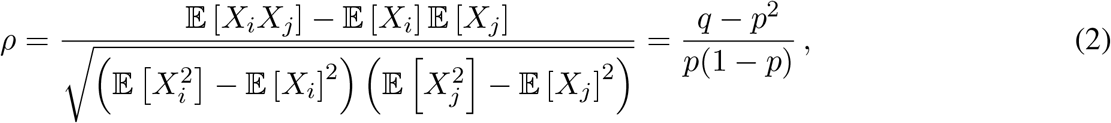

so that we have *q* = *p*^2^ + *ρp*(1 − *p*). In this context, the *χ*-metric can be computed as

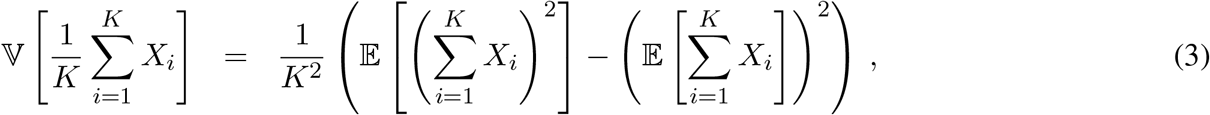

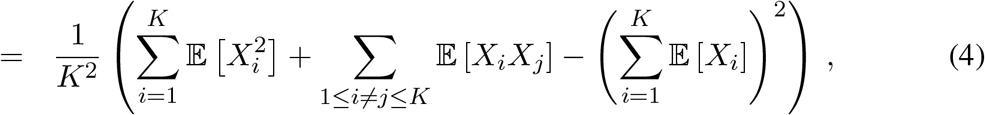

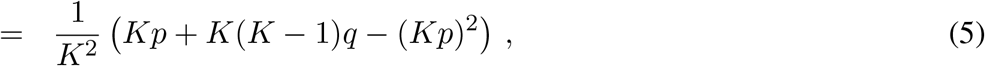

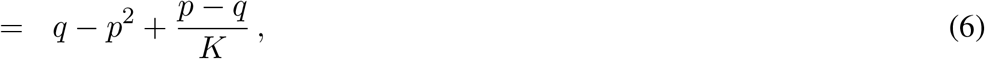

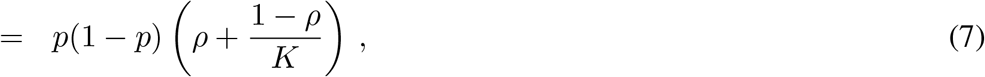

where the last line is obtained by expressing *q* in term of the spike correlation *ρ*. For spiking data, we commonly have *p* = *r*Δ*t* ≪ 1, so that *p*(1 − *p*) ≃ *p* = *r*Δ*t*, leading to the rate normalized *χ*-metric

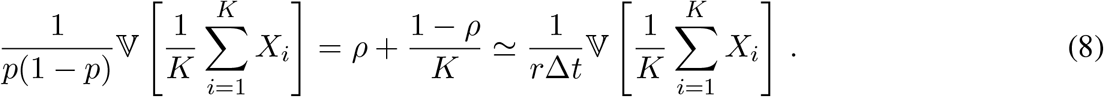

The above relation demonstrates that for the Bernoulli homogeneous case, studying the (1*/K*)-dependence of the *χ*-metric allows one to rigorously infer the shared spiking correlation.

##### 1.1.2 Binomial counts

Assuming no temporal correlations, the above result generalizes to the spike counts observed in many time bins. Specifically, for an integer bin size *M*, we have 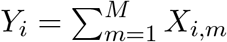 where 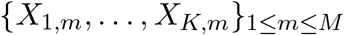, are independent samples across time. This corresponds to taking:

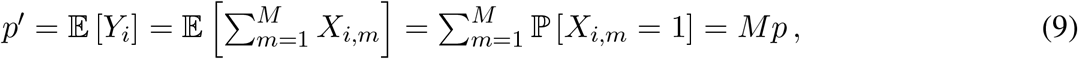

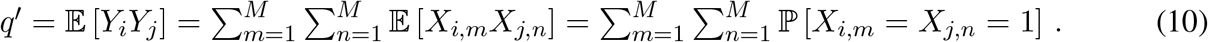

Separating the contributions of coincidental time bins and noncoincidental (independent) time bins yields

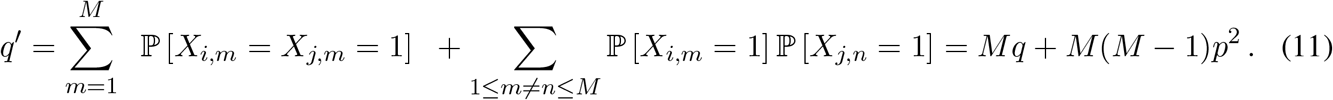

Similarly, we have

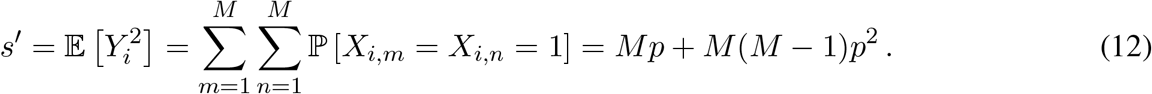

Using the same calculation as for the Bernoulli case, this allows one to evaluate the new normalized *χ*-metric

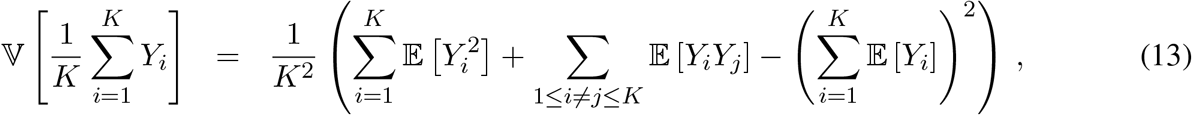

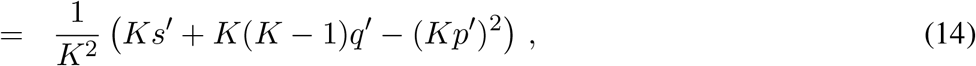

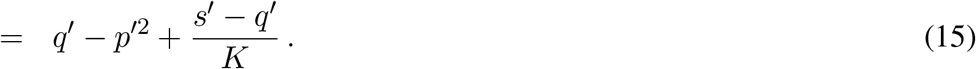

Expressing *p*^′^, *q*^′^, and *s*^′^ in terms of *p* and *q*, we get

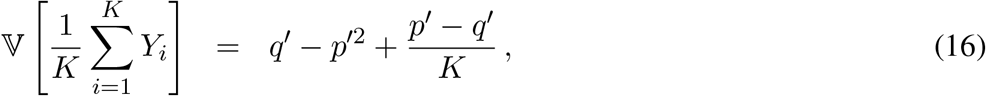

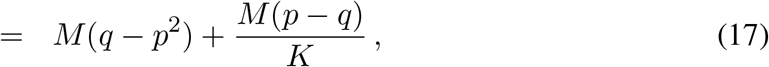

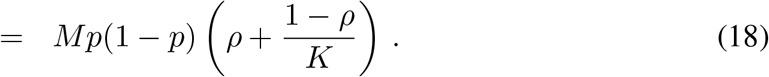

Thus, after proper rate normalization, larger bin sizes do not affect the (1*/K*)-dependence.

##### 1.1.3 Poisson counts

From there, another generalization is to consider Poisson distributed spike counts, given a common spiking rate *r*, this classically corresponds to considering

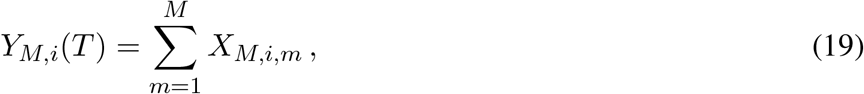

in the limit *M* → ∞, where {*X*_*M*,1,*m*_, …, *X*_*M,K,m*_}_1≤*m*≤*M*_ is the spiking activity in a time bin of size *T/M* so that we have 𝔼 [*X*_*M,i,m*_] = *p*_*M*_ = *rT/M* = *p/M*. Assuming that the pairwise correlation persists at small timescale is equivalent to assuming that the scaling *q*_*M*_ = 𝔼 [*X*_*M,i,m*_*X*_*M,j,m*_] = *q/M* holds as well. This scaling corresponds to an instantaneous form of synchrony, which we discuss in more detailed in Section 2.1. This instantaneous form of synchrony follows from the fact that under the considered scaling, one obtains limit Poisson processes *N*_*i*_(*T*) = lim_*M*→∞_ *Y*_*M,i*_ which can jump synchronously. One can derive the corresponding limit *χ*-metric by first observing that

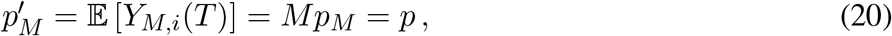

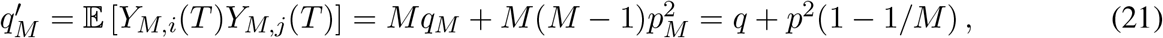

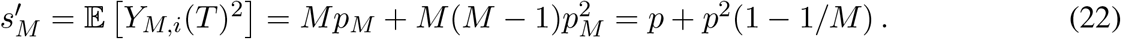

This leads to the limit pairwise spiking correlation

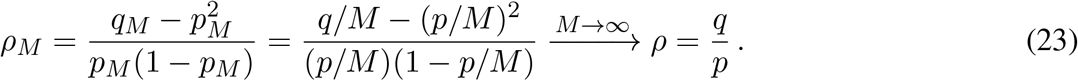

Using the above results, we have

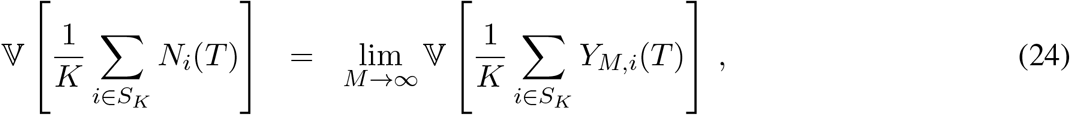

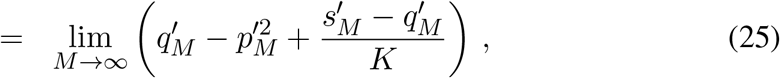

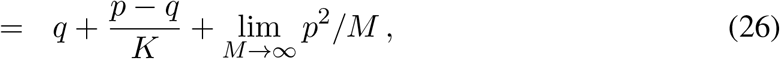

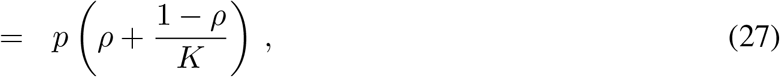

where the last equality follows from substituting *q* = *ρp*. This leads to the instantaneous Poisson version of the *χ*-metric

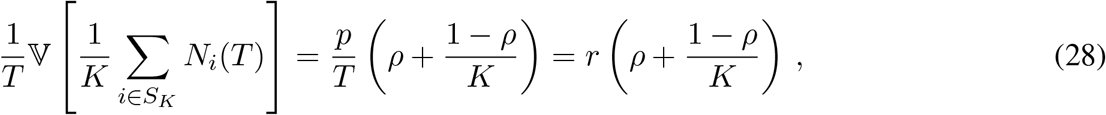

which exhibits a similar (1*/K*)-dependence as for the binomial case.

#### 1.2 Population heterogeneity

##### 1.2.1 Discrete-time model

Let us consider the more realistic case of heterogeneous spiking rates and heterogeneous correlations, which we parametrized via

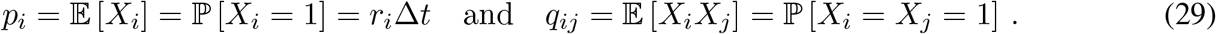

For an integer bin size *M*, the quantities of interest are the spike counts 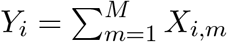, where the vectors { *X*_1,*m*_, …, *X*_*K,m*_ } _1≤*m*≤*M*_, are independent samples across times. Similarly to the homogeneous case, we introduce the following useful quantities:

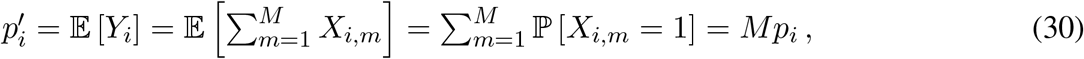

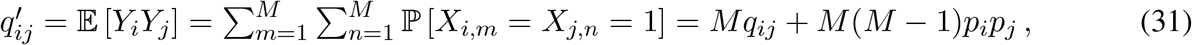

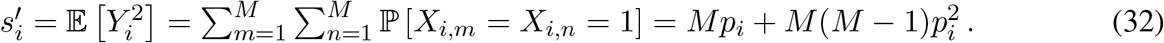

It is also convenient to compute the binned covariance as

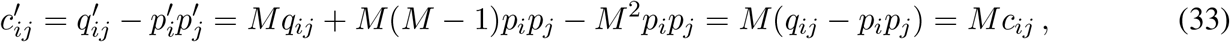

where *c*_*ij*_ is the single-bin covariance. Taking into account heterogeneities causes the *χ*-metric to depend on the set of *K* neurons under consideration, which we denote by *S*_*K*_ ⊂ {1, …, *K*}. With this in mind, we can evaluate

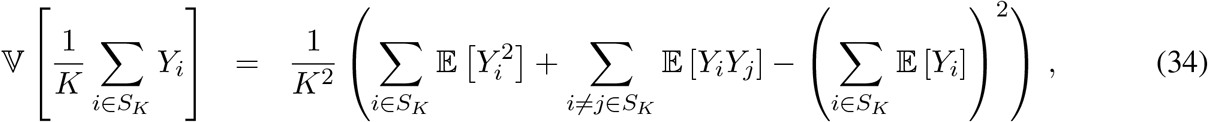

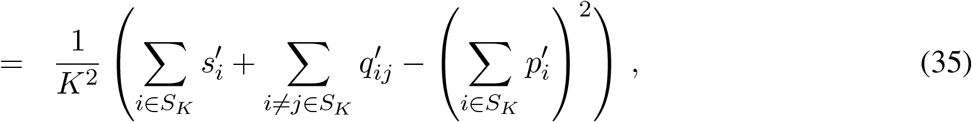

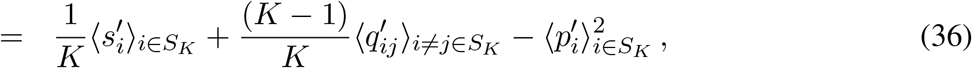

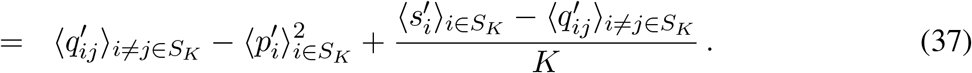

Due to heterogeneities, there are actually (1*/K*)-correction terms in the inhomogeneous term:

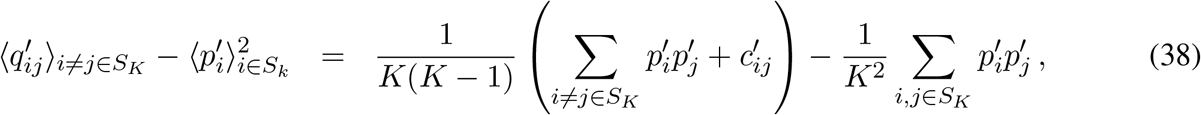

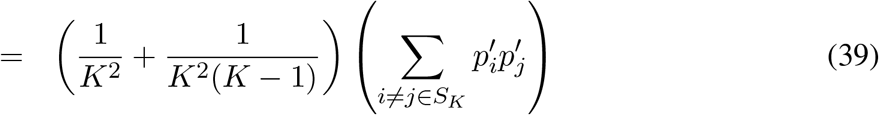

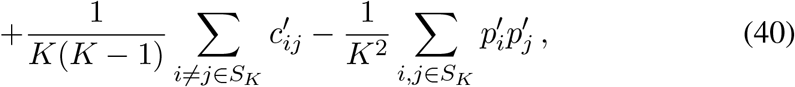

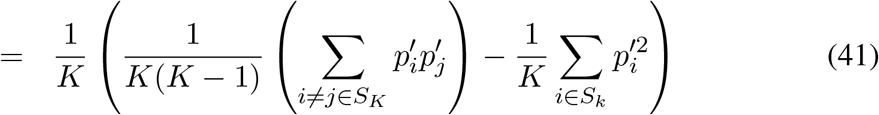

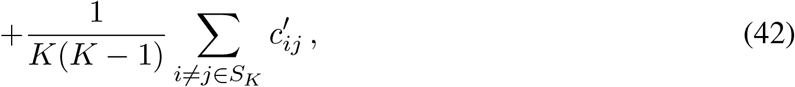

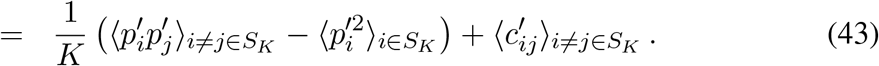

Thus, we have the following overall (1*/K*)-dependence for the *χ*-metric

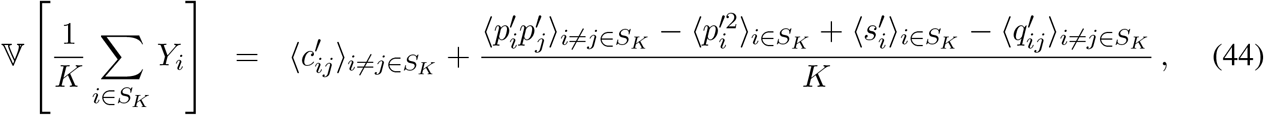

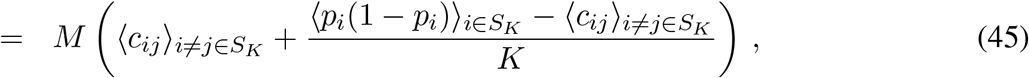

where the last equality follows from observing that

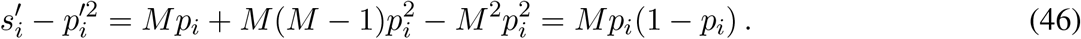

In the presence of population heterogeneity, one thus obtain the (1*/K*)-dependence for the *χ*-metric by merely performing the population averages of the involved statistics.

##### 1.2.2 Continuous-time model

Obtaining the continuous-time model amounts to taking the Poissonian limit of the discrete model, which involves considering time bins of duration Δ*t* = *T/M* with *M* → ∞. Taking such a Poissonian limit yields instantaneously synchronous Poisson processes *N*_*i*_, whereby each process register the spiking activity of neuron *i*. Moreover, one can check that in the Poissonian limit, the *χ*-metric satisfies

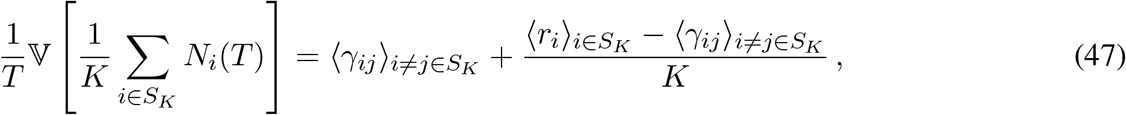

where the instantaneous covariance coefficients *γ*_*ij*_ = *Mc*_*ij*_*/T* = *c*_*ij*_*/*Δ*t* satisfy ℂ[*dN*_*i*_(*t*)*dN*_*j*_(*t*)] = 𝔼 [*dN*_*i*_(*t*)*dN*_*j*_(*t*)] = *γ*_*ij*_*dt*. Finally, the *χ*-metric can be written as

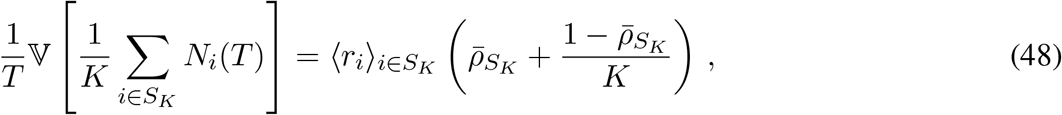

where we have defined the correlation-like coefficient 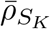 as the ratio of the average instantaneous co-variance with respect to the average instantaneous variance: 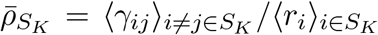. Note that the latter quantity is distinct from the average correlation. For instance, consider the case of a constant pair-wise spiking correlation *ρ* so that we have 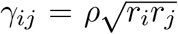. Then, 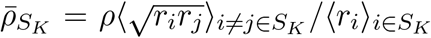, which differs from *ρ* in general. Note also that we have assumed the set of neurons *S*_*K*_ fixed throughout the calculation. Numerical estimation of *χ*-metric actually benefits from evaluating the (1*/K*)-dependence via sample-average variance estimate obtained by randomly sampling *S*_*K*_ for intermediate *K*. This corresponds to altering the results by performing an additional average over *S*_*K*_, but should not change the essence of the result.

#### 1.3 Temporal correlations

##### 1.3.1 Discrete-time model

Finally, we consider a spiking correlation model that also includes temporal correlations. For simplicity, we consider that these temporal correlations are homogeneous in time and identical across neuronal pairs. For a Bernoulli model, this corresponds to considering for all 1 ≤ *i* ≤ *K*, 1 ≤ *m* ≤ *M*

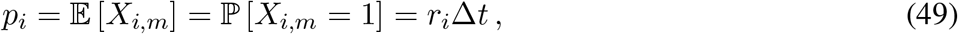

and for all 1 ≤ *i, j* ≤ *K*, 1 ≤ *m, n* ≤ *M* with (*i, m*) ≠ (*j, m*)

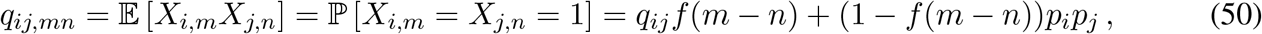

where the function *f* quantifies temporal correlations and is such that 0 ≤ *f* ≤ 1 with *f* (0) = 1. Observe that by definition *q*_*ii,mm*_ = 𝔼 [*X*_*i,m*_*X*_*i,m*_] = 𝔼 [*X*_*i,m*_] = *p*_*i*_. A typical example of such functions that involves a single time scale *τ* is given by *f* (*n*) = exp (−| *n*| Δ*t/τ*), where Δ*t* denotes the duration of a bin. Similarly to the temporally independent case, we introduce the following useful quantities:

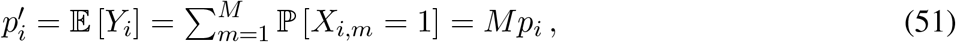

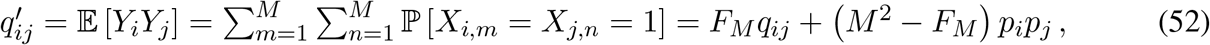

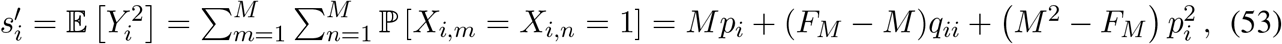

where we have defined

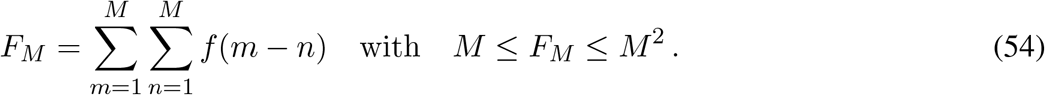

Observe that using (50), *F*_*M*_ can be interpreted as

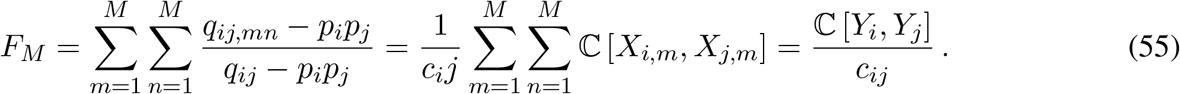

where *c*_*ij*_ is the single-bin covariance. It is also convenient to compute the binned covariance as

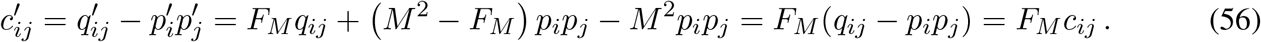

Following the same calculations as for the case without temporal correlations, we find overall (1*/K*)-dependence for the *χ*-metric to be

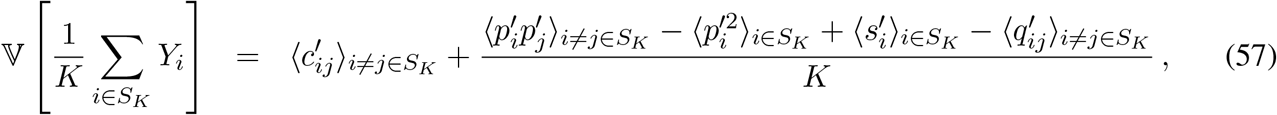

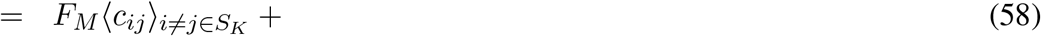

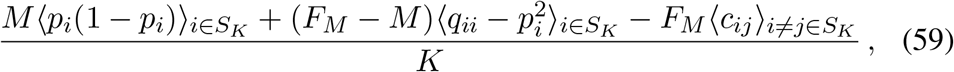

where the last equality follows from observing that

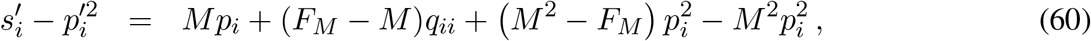

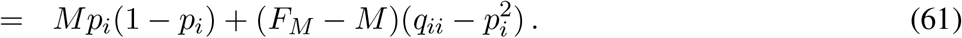

Thus, we still obtain the characteristic (1*/K*)-dependence of the *χ*-metric observed in the absence of temporal correlation, albeit with a scaling factor *F*_*M*_ that depends on the duration of the bin as well as on the time scale of the correlations.

##### 1.3.2 Continuous-time model

As before, obtaining the continuous-time model amounts to taking the Poissonian limit of the discrete model, which involves considering time bins of duration Δ*t* = *T/M* with *M* → ∞. By contrast with the temporally independent case, the corresponding limit counting processes specified by *N*_*i*_(*T*) = lim_*M*→∞_ *Y*_*i*_ are Poisson processes with stochastic rates, which are commonly referred to as doubly-stochastic processes. Moreover, in the presence of temporal correlations, one expects that *F*_*M*_ scales as *M* ^2^. For instance, one can check that for *f* (*n*) = exp (−| *n*| *T/Mτ*), we have

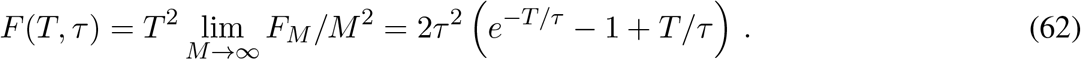

This is by contrast with the temporally independent case for which *F*_*M*_ ∼ *M* and suggests that we consider a different form of scaling for the parameters *q*_*ij*_, which are assumed to satisfy *q*_*ij*_ ∼ Δ*t*∼ 1*/M* when modeling instantaneous synchrony. Clearly, one has to assume that *q*_*ij*_∼ Δ*t*^2^ ∼1*/M* ^2^, a scaling that is naturally achieved for doubly stochastic models. To see why, let us consider that neuron *i* and neuron *j* spike according to two conditionally independent Poisson processes with correlated stochastic rates denoted by *Z*_*i*_(*t*) and *Z*_*j*_(*t*), respectively. Given that we have 𝔼 [*Z*_*i*_] = *r*_*i*_, and denoting *Z*_*i,m*_ = *Z*_*i*_(*m*Δ*t*) and *Z*_*j,n*_ = *Z*_*j*_(*n*Δ*t*), we model *X*_*i,m*_ and *X*_*j,n*_ as conditional independent Bernoulli variables with parameters *Z*_*i,m*_Δ*t* and *Z*_*j,n*_Δ*t*, respectively. Then, one can check that as expected, we have

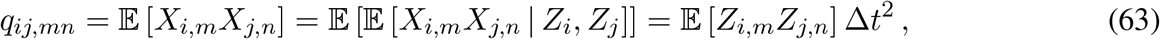

The above scaling, achieved for doubly-stochastic models, corresponds to a more realistic form of synchrony that does not assume instantaneously synchronous spiking and which we discuss in Section 2.2. To connect the doubly-stochastic model with our discrete model, observe that

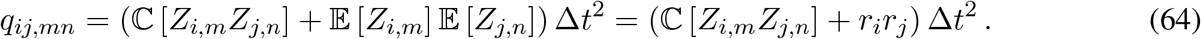

Then, comparing the above expression with (50) shows that doubly-stochastic models correspond to choosing *f* such that

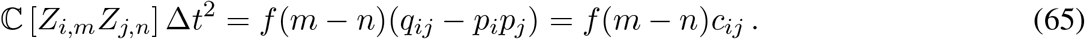

In particular, we have the limit behaviors

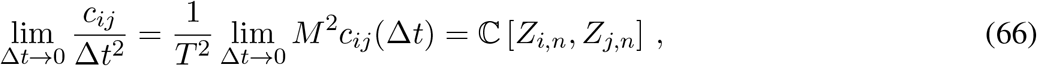

In turn, denoting *ζ*_*ij*_ = ℂ[*Z*_*i,n*_, *Z*_*j,n*_], we obtain the Poissonian limit for the *χ*-metric as

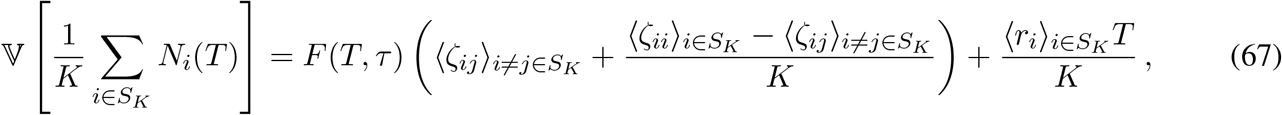

which again exhibits the characteristic (1*/K*)-dependence but where the scaling factor *T* ^2^*F* (*T, τ*) depends on the bin duration *T* and the correlation time scale *τ*. Tellingly, one can interpret the scaling factor *T* ^2^*F* (*T, τ*) in term of the doubly-stochastic processes *N*_*i*_(*t*) by observing that

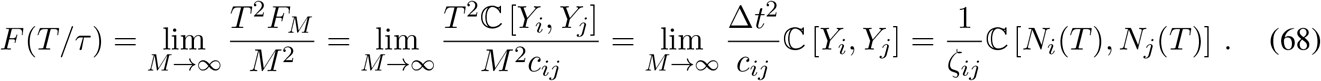

One can further deduce the expression of *F* (*T, τ*) from the rate crosscorrelations ℂ [*Z*_*i*_(*s*), *Z*_*j*_(*t*)] (see Section 2.2.3) by evaluating

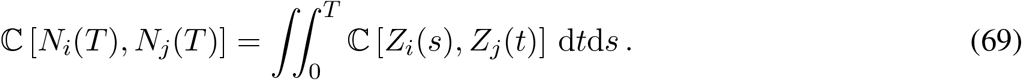

For instance, one can check that the limit expression (62) consistently corresponds to choosing ℂ [*Z*_*i*_(*s*), *Z*_*j*_(*t*)] = *ζ*_*ij*_*e*^−| *t*−*s*| */τ*^. Finally, the above remarks allows one to check that as expected, expression (67) for the *χ*-metric is indeed equivalent to

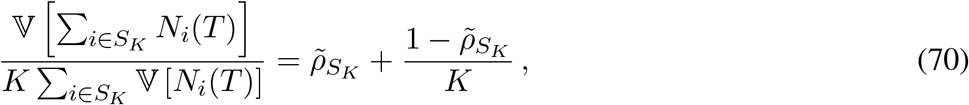

where we have defined the correlation-like coefficient 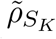 as the ratio of the average covariance with respect to the average variance: 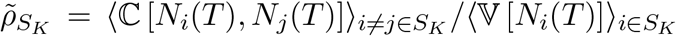. Note that as for 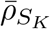, the quantity 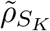 is again distinct from the average correlation in the presence of population heterogeneities. In the presence of temporal correlation, the main difference is that the normalization of the *χ*-metric must involved the variance rather than the rate alone as: 𝕍 [*N*_*i*_(*T*)] = 𝔼 [*Z*] *T* + 𝕍 [*Z*] *T* ^2^ = *r*_*i*_*T* + *ζ*_*ii*_*T* ^2^ *> r*_*i*_*T* (see Section 2.2.3).

### 2 Synchronous spiking input models

In this section, we discuss spiking population models exhibiting two distinct form of synchrony. The first class exhibits instantaneous synchrony, whereby neurons’ spiking activity are governed by Poisson processes which can jump simultaneously. The second class exhibits a looser but more realistic form of synchrony, whereby neurons are governed by Poisson processes with fluctuating, correlated rates. The focus is one deriving parametric forms for the correlation coefficients and for the Fano factors of the neuronal spike counts.

#### 2.1 Instantaneous synchrony

For simplicity, we consider exchangeable models for which neurons are assumed to be pooled from a large (infinite-size) reservoir of identically acting neurons. In this context, instantaneous-synchrony models correspond to assuming that synchrony arises from an independently fluctuating mean spiking count across time bins. When this probability is high, neurons tend to coactivate; when this probability is low, neurons tend to remain collectively silent. Such mean count fluctuations thereby lead to overall spiking synchrony.

##### 2.1.1 The Poisson-gamma model

The Poisson-gamma model is defined by assuming that the common mean spiking count of *K* exchangeable Poisson neurons follows a gamma distribution *µ*_*α,β*_. Specifically, given a time bin of duration Δ*t*, we consider *µ*_*α,β*_ = Gamma(*α, β*) with parameter *α/β* = *r*Δ*t* for some *r >* 0 and *β >* 0. Then, the spike-count vector *N*_1_, …, *N*_*K*_ is given by

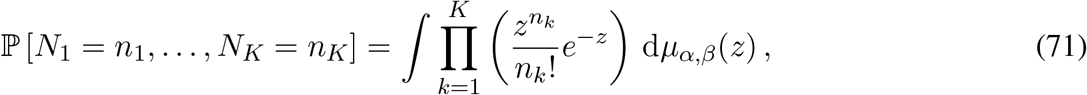

For exchangeable neurons, the spike-count correlation is defined as

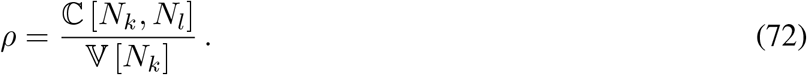

For an exchangeable Poisson model with directing mean count probability *µ*_*α,β*_, we have

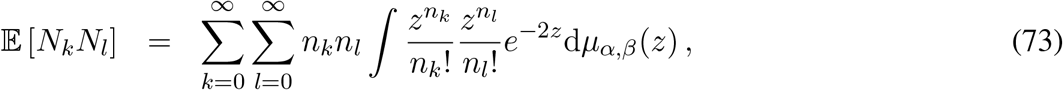

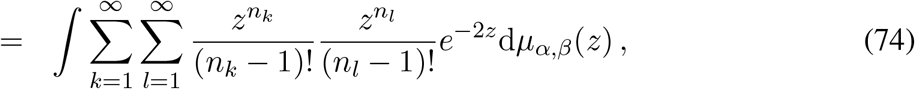

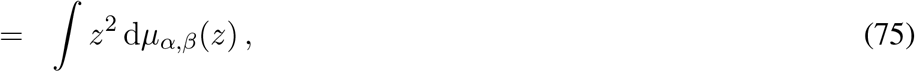

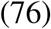

so that we have ℂ [*N*_*k*_, *N*_*l*_] = 𝔼 [*N*_*k*_*N*_*l*_] − 𝔼 [*N*_*k*_]^2^ = 𝔼 [*Z*^2^] − 𝔼 [*Z*]^2^ = 𝕍 [*Z*]. At the same time, we also have

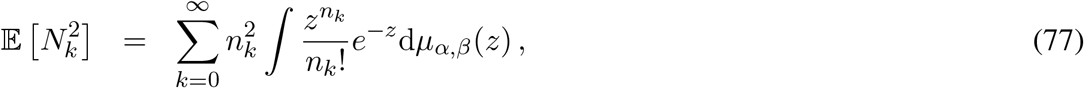

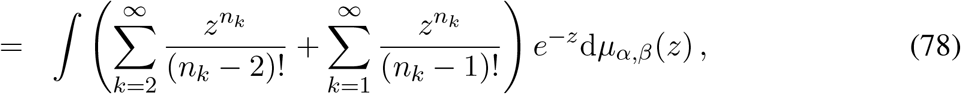

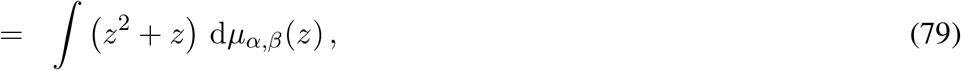

so that we have 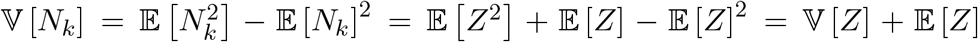. Thus the instantaneous spike-count correlation is specified for an exchangeable Poisson model as

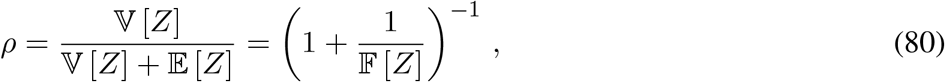

where 𝔽 [*Z*] denotes the Fano factor of the underlying mean count *Z*. This can further be specified for the Poisson-gamma model by using the fact that 1*/*𝔽 [*Z*] = *β* to obtain *ρ* = 1*/*(1 + *β*), so that *β* parametrizes spike-count correlations.

##### 2.1.2 Independent Poisson-gamma process

Given a sequence of *M* time bin of duration Δ*t*, let us consider that the spiking activity in each time bin follows an independent Poisson-gamma model. For a *K*-neuron model, this means that the probability of the time-indexed population vector {*N*_*k,m*_}, 1 ≤ *k* ≤ *K*, 1 ≤ *m* ≤ *M*, with 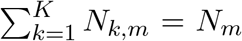, is specified by

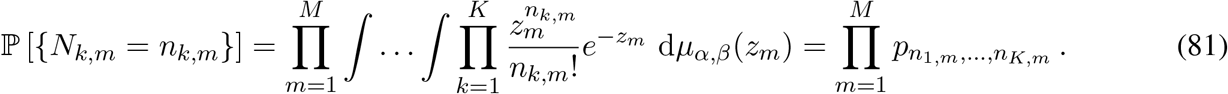

As we consider exchangeable models for which every inputs play the same role, it is actually enough to track the total spike count 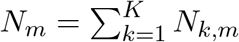 in each time bin. The corresponding time-indexed population vector {*N*_*m*_}, 1 ≤ *m* ≤ *M*, satisfies

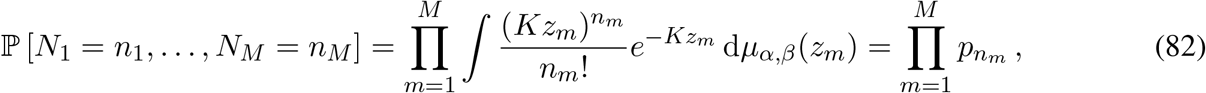

A nice feature of the independent Poisson-gamma process is that it enjoys divisibility in the sense that the functional form of its probability law is stable under dividing and merging bins. For fixed correlation *ρ*, i.e., for fixed *β*, this follows from the additivity of Poisson and gamma random variables. For instance, one can check that upon merging *M* bins, one has

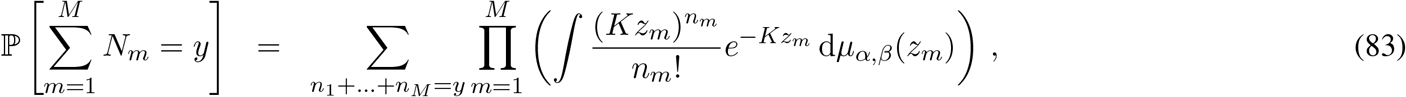

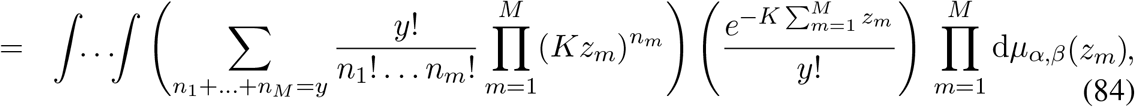

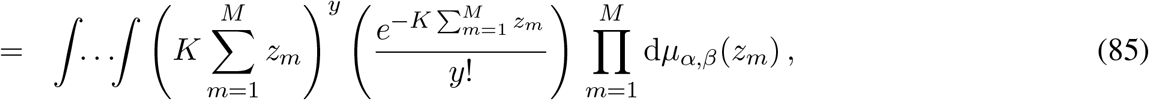

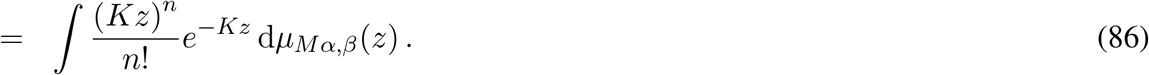

The above observation implies that the spike-count correlation *ρ* is independent of the bin size *M* Δ*t*. Similarly, the cell-specific Fano factor is also independent of the bin size *M* Δ*t* as one can check that

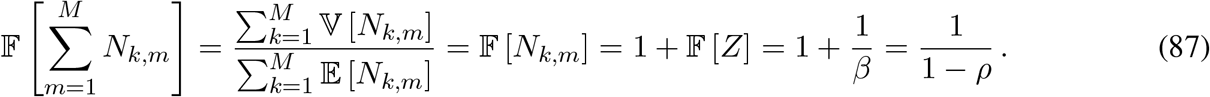

Accordingly, the spike-count correlation *ρ* is also independent of the number of bins *M* :

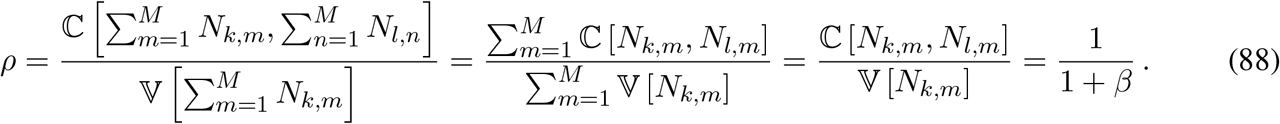

Finally, denoting by *Z*_*M*_ a gamma random variable with parameter (*Mα, β*), we can check that the population Fano factor is given by

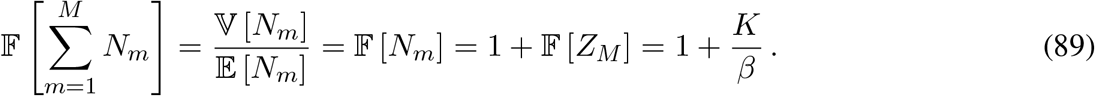

##### 2.1.3 Continuous-time limit for the independent Poisson-gamma process

The continuous-time limit is obtained by considering the discrete spiking model for time bins of duration Δ*t* = *T/M* with *M* → ∞. By exchangeability of the discrete model, it is enough to keep track of time-indexed the total spike count 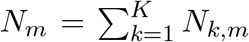 in each time bins. Given our additional assumption that *N*_1_, …, *N*_*m*_ are i.i.d., let us denote by *N* a generic total spike count. By additivity of Poisson random variables, the distribution of *Y* over {0, 1, …, *K*} is given by

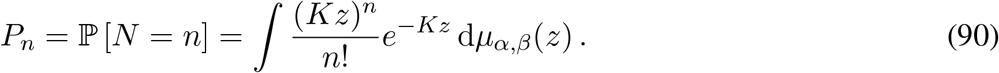

Evaluating the above integral yields the distribution of the counting variable *N* as

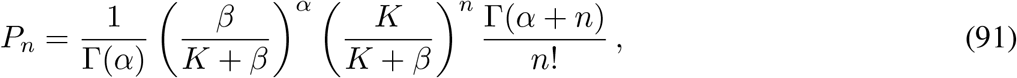

which is always summable as

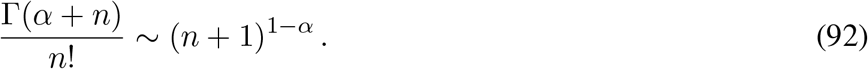

It is then convenient to think of the discrete process 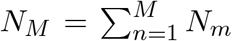 as an integer-valued random walk with i.i.d. positive jumps, denoted by *k*. By construction, these jumps have distribution *P*_*k*_*/*(1 − *P*_0_) on ℕ ^*⋆*^. In the discrete model, the mean spiking count is given by 𝔼 [*Z*] = *α/β* = *rT/M*, so that one can achieve the continuous-time limit *M*→ ∞ by considering *α* → 0. In this continuous-time limit, by independence across time bins, one can then show that the random walk *Y*_*M*_ tends to a compound Poisson process *Y* (*t*) such that for all *t >* 0

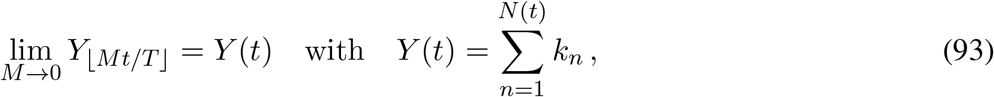

where *N* is a Poisson process with rate *b* and *k*_*n*_ are i.i.d. with common distribution

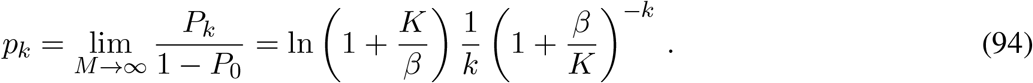

The rate *b* of the driving Poisson process *N* is specified by the conservation of the overall spiking rate, which imposes that *b*𝔼 [*k*] = *Kr* so that:

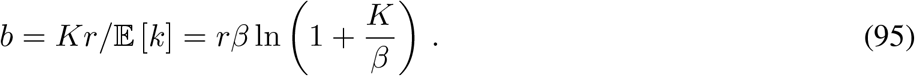

Observe that continuous-time limit *α* → 0 leaves the correlation coefficients and the Fano factor unchanged as these only depend on the parameter *β* and *K*. In turn, these parameters entirely specify the common jump distribution *p*_*k*_, which explain the emergence of perfect synchrony in the presence of nonzero spiking correlations. In the absence of correlations (*ρ* = 0 and *β* = ∞), synapses spike asynchronously so that only one synapse activates at a time: *k* = 1 with probability one, i.e., *p*_1_ = 1. In the presence of correlations (*ρ >* 0 and *β <*∞), synapses act synchronously so that many synapses activate at the same time: *k >* 1 with nonzero probability, i.e., *p*_1_ *<* 1.

#### 2.2 Loose synchrony

For simplicity, we still consider exchangeable models for which neurons are assumed to be pooled from a large (infinite-size) reservoir of identically acting neurons. In this context, we model a loose form of spiking synchrony by assuming that this synchrony arises from a collective fluctuating spiking rate (as opposed to a mean spiking count) across time bins. Such models belong to the class of doubly-stochastic models and allow for the introduction of temporal correlations across time bins. This is by contrast with instantaneous models, which rely on the assumption of temporal independence. Synchrony is established by stochastically alternating periods of elevated and depleted spiking rate. This form of synchrony is more realistic for not relying on exactly synchronous spiking across neurons.

##### 2.2.1 Basic facts about the CIR process

We model synchrony with finite temporal correlation by assuming that the underlying firing rate follows a Cox-Ingersoll-Ross (CIR) dynamics. Specifically, we consider that the joint firing rate *Z*_*t*_ of an exchangeable sequence of *K* Poisson neurons follows the stochastic equation

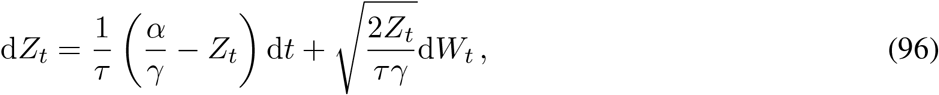

where *W*_*t*_ denotes the canonical Brownian motion. In the above equation, *α* is a positive dimensionless parameter, *γ* is a positive time scale, both of which are to be discussed later, and *τ* is the correlation time of the dynamics. Denoting *ζ* = *e*^−*t/τ*^, the CIR Markov transition kernel is known analytically as

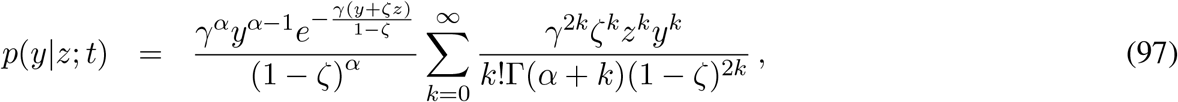

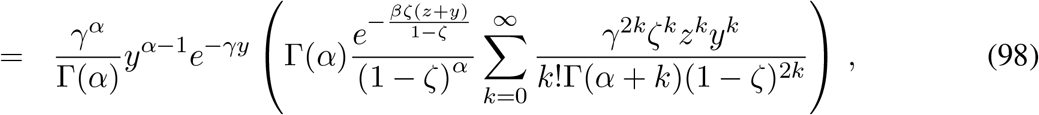

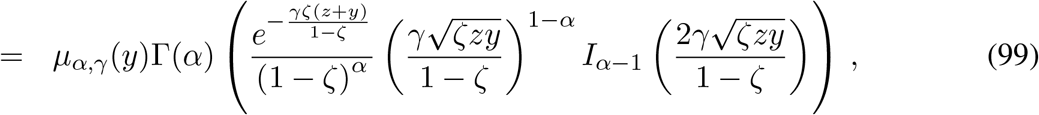

where *I*_*α*−1_ denotes the modified Bessel function of the first kind with parameter *α* −1. Moreover, the moment-generating function of the transition kernel is also known as

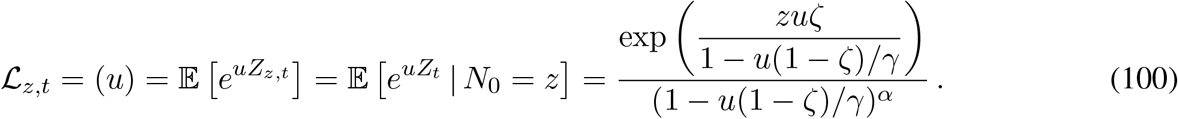

From there, one can compute that

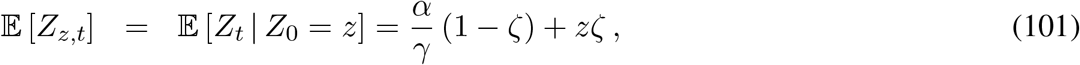

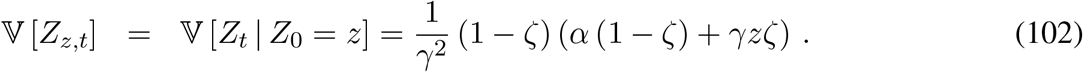

The above CIR dynamics admits a stationary distribution given as the gamma distribution

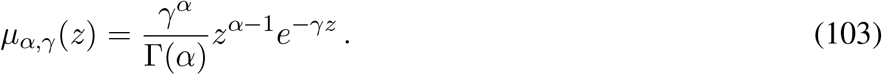

Considering a CIR process with initial condition *N*_0_ distributed as *µ*_*α,γ*_ specifies the stationary CIR process, simply denoted as *Z*_*t*_, for which we have

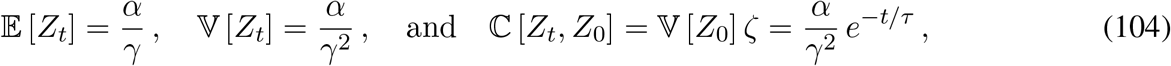

confirming that *τ* as a correlation time constant, whereas *γ* can be interpreted as the inverse stationary Fano factor *γ* = 𝔼 [*Z*_*t*_] */*𝕍 [*Z*_*t*_] = 1*/*𝔽 [*Z*_*t*_]. The later quantity must have unit of a time, since *Z*_*t*_ will be interpreted as a rate in the following (as opposed to being spike counts as for the Poisson-gamma model).

##### 2.2.2 The Poisson-CIR process model

Given a sequence of *M* time bins of duration Δ*t*, let us consider that the common spiking rate in each time bin follows a stationary CIR rate process *Z*_*t*_. Then, the probability of time-dependent spike-count trajectory {*N*_1_, … *N*_*M*_ } is given by

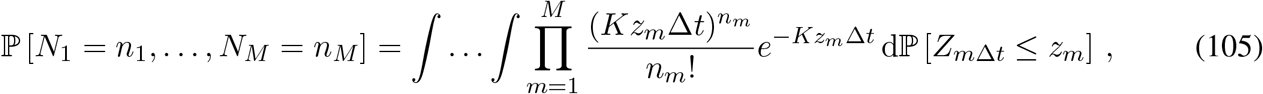

whereas the population vector {*N*_*k,m*_}, 1 ≤ *k* ≤ *K*, 1 ≤ *m* ≤ *M*, with 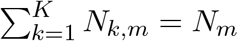, satisfies

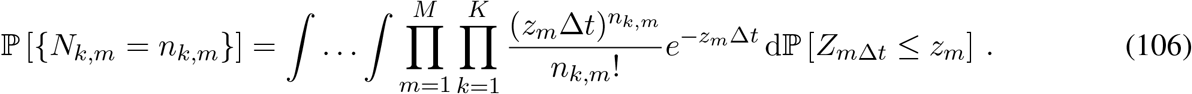

In order to compute the spike-count correlation over bins of size *M* Δ*t*, we first consider the covariance

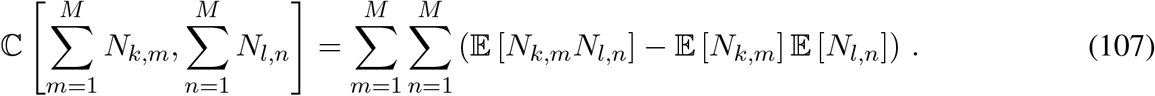

For (*k, m*) ≠ (*l, n*), the individual covariance terms above evaluate to

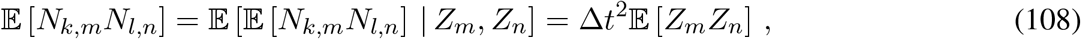

whereas for (*k, m*) = (*l, n*), we have

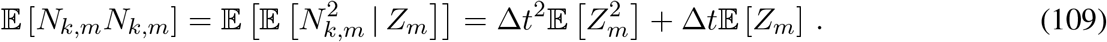

Thus, the spike-count correlation *ρ*_*kl*_ between neurons *k* and *l* is given in term of the underlying CIR process *Z* as

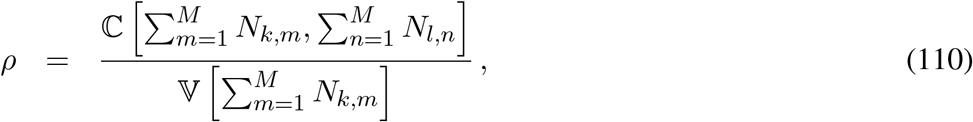

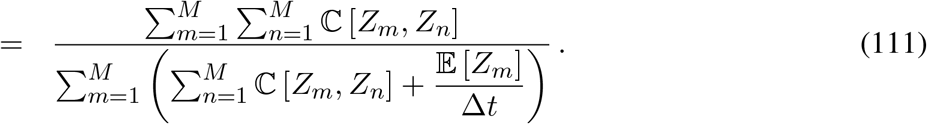

Exploiting the temporal correlation structure of the CIR process, we further obtain

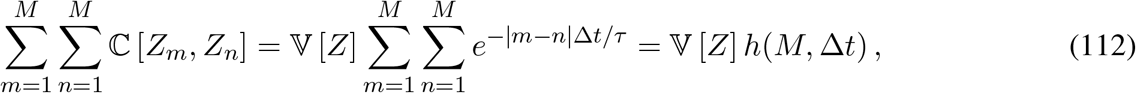

where the auxiliary function *h* captures the bin-size dependence via

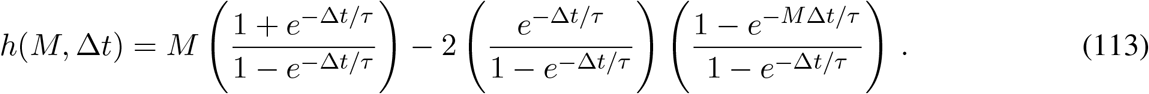

One can check that at fixed Δ*t*, the function *f* is monotonically increasing with

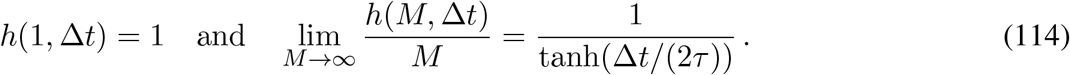

This leads to the bin-size-dependent spike-count correlation

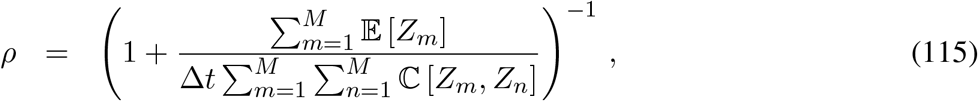

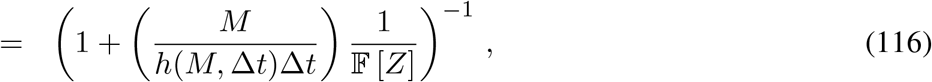

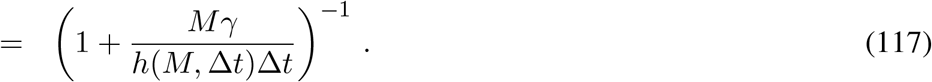

The above results also allow us to compute the Fano factor

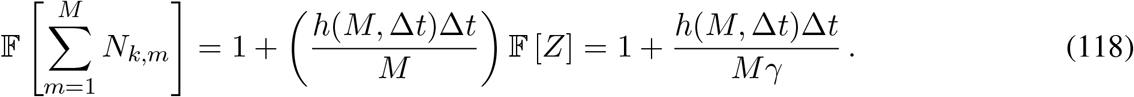

Similar calculations about the total spiking count yield

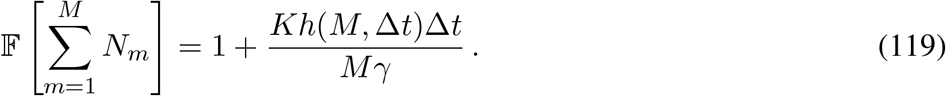

Observe that the above results differs from those obtained from the Poisson-gamma model in two ways: First, the presence of temporal correlations implies the occurrence of the multiplicative term *M/h*(*M*, Δ*t*), which consistently tends to one when the correlation time vanishes, i.e., *M/h*(*M*, Δ*t*) →1 when *τ* → 0. Second, when *τ* → 0, the spike-count correlation for an homogeneous population reads *ρ* = 1*/*(1 + *γ/*Δ*t*), indicating that *γ/*Δ*t* plays the role of the dimensionless parameter *β* in the Poisson-gamma model. In particular, this shows that one expect instantaneous spike-count correlations to vanish in the limit of infinitesimal bin size: Δ*t* → 0.

##### 2.2.3 Continuous-time limit for the CIR-process model

In the continuous-time limit, the discrete spiking model naturally converges toward a doubly-stochastic process, whereby all neurons share the common stochastic CIR rate *Z*(*t*). One can derive the associated spiking correlation and Fano factor at time scale *T* by setting Δ*t* = *T/M* and taking the limit *M* → ∞. One then finds the limiting behavior

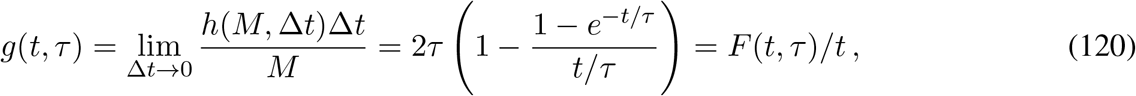

where the newly introduced function *g* is increasing with respect to *t* over ℝ^+^ with *g*(0, *τ*) = 0, *∂*_*t*_*g*(0, *τ*) = 1, and lim_*t*→∞_ *g*(*t, τ*) = 2*τ*. Thus, in the continuous-time limit, one can evaluate the spike-count correlation between inputs *k* and *l* and for bin size *T* as

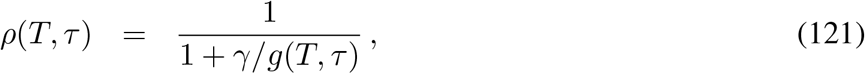

as well as the associated Fano factors for the individual spike count *N*_*k*_ and for the total spike count 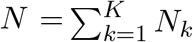:

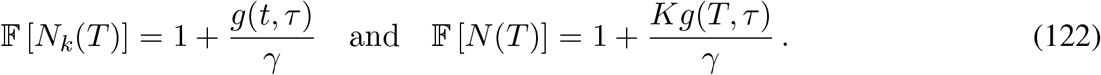

These results can be directly derived from considering the doubly-stochastic processes *N*_*k*_, 1 ≤ *k* ≤ *K* governed by the common rate *Z*(*t*). Specifically, for such processes, one has the infinitesimal covariance

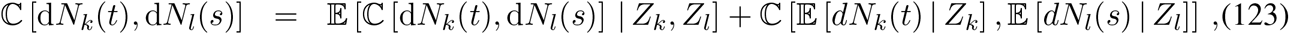

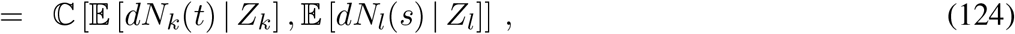

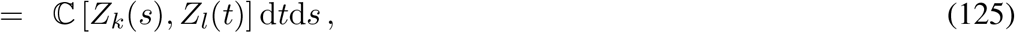

and the infinitesimal variance

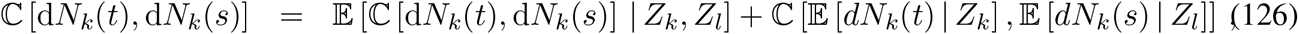

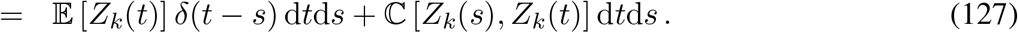

Assuming that 𝔼 [*Z*_*k*_(*t*)] = *α/γ* and ℂ [*Z*_*k*_(*s*), *Z*_*l*_(*t*)] = (*α/γ*^2^) *e*^| *t*−*s*| */τ*^ and using that

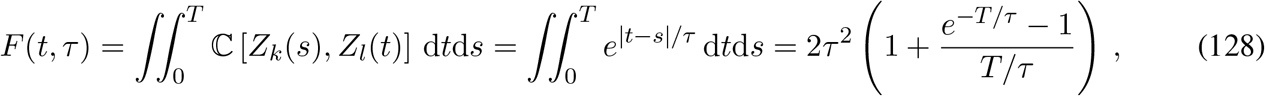

we find that

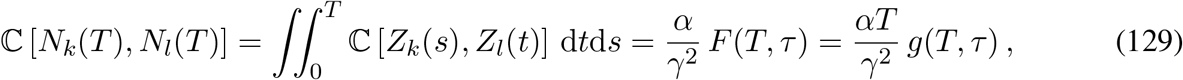

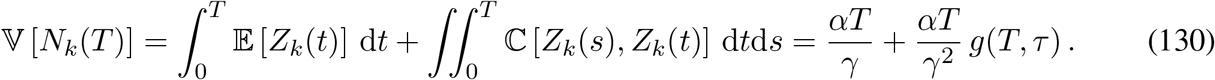

As expected, this leads to

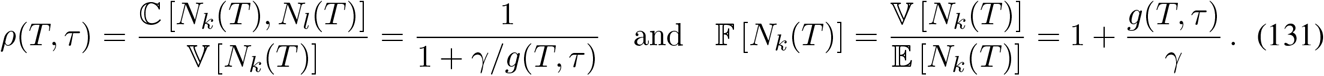

### 3 Shot-noise model for conductances

In this section, we discuss various conductance shot-noise models derived from considering synaptic inputs with synchrony. The focus is on deriving a parametric form for a measurable quantity, called *Q*, that can serve to assess the degree of synchrony compatible with a conductance measurement.

#### 3.1 Asynchronous input drive

We consider that the activation of *K* asynchronous synapses is governed by independent Poisson processes with rate *r*. We further assume that synaptic activations elicit conductance changes of typical amplitude *A* with synaptic time constant *τ*_s_. Accordingly, the overall conductance is modeled as a shot noise and in the stationary regime, we have

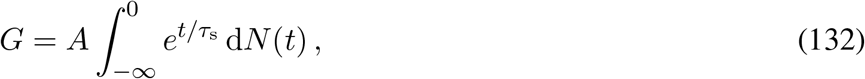

where the governing Poisson process *N* has overall input rate *Kr*. The mean stationary conductance can be computed as

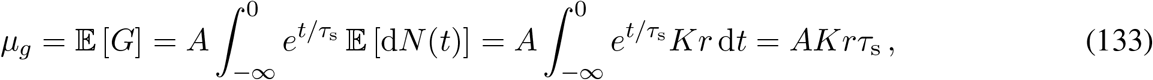

whereas the stationary conductance variance is given by

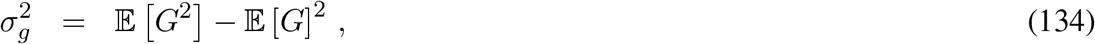

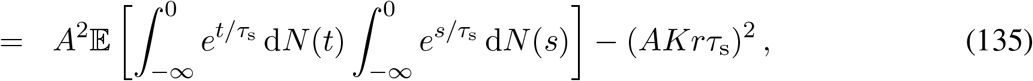

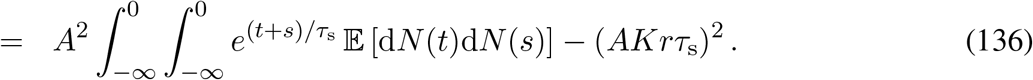

Using that for a Poisson process with rate *Kr*, we have 𝔼 [d*N* (*t*)d*N* (*s*)] = [(*Kr*)^2^ + *Krδ*(*t s*)] d*t*d*s*, we obtain:

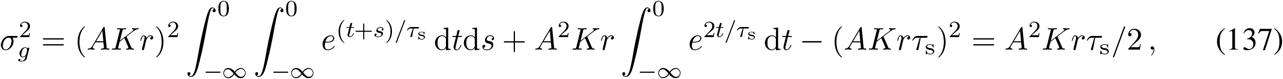

which implies that we can form the amplitude-independent quantity

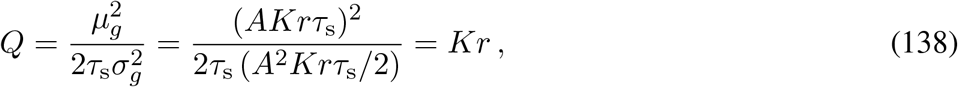

which happens to e equal to the overall input rate *Kr* in the asynchronous regime.

#### 3.2 Input drive with instantaneous synchrony

We simulate instantaneous synchrony by allowing for several synaptic inputs to activate at the exact same time. This corresponds to modeling the synaptic drive via a compound Poisson process rather than a Poisson process. In our case, we consider a compound Poisson process *Y* defined as

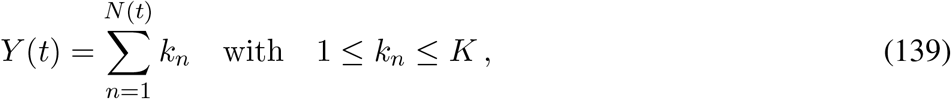

where *k*_*n*_ are independent, identically distributed, integer-valued jumps in {1, …, *K*}. The jumps *k*_*n*_, which we will refer to as *k* when it is not ambiguous, represents the possibly fluctuating numbers of coactivating synaptic inputs. Specifically, one can show that the spiking correlation coefficient *ρ* is independent of the bin size and satisfies

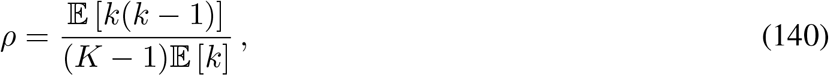

where 𝔼 [·] denotes expectation with respect to the jump distribution *p*_*k*_. As usual, the rate of the governing Poisson process *N*, which we denote *b*, is chosen so that the overall input rate is preserved, independent of synchrony. This implies that one must choose *b* such that *b*𝔼 [*k*] = *Kr*.

Given these preliminary remarks, the conductance process resulting from synaptic inputs with instantaneous synchrony is specified as the compound-Poisson-process shot noise

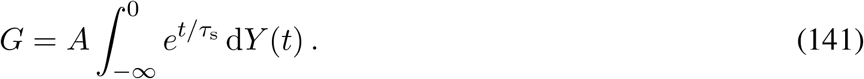

For a fixed overall input rate, the stationary mean conductance is independent of synchrony as

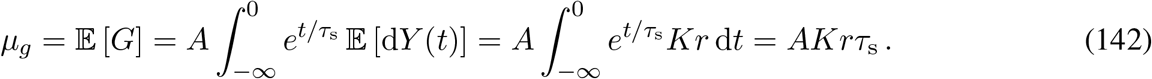

In turn, the stationary conductance variance in the presence of synchrony can be evaluated as

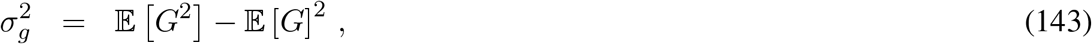

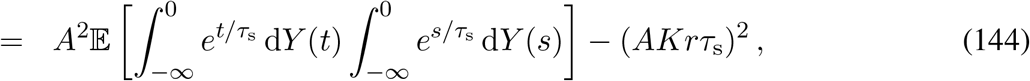

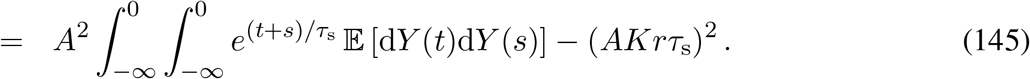

For a compound Poisson process *Y* with jumps *k* and rate *b*, we have 𝔼 [d*Y* (*t*)d*Y* (*s*)] = [(*b*𝔼 [*k*])^2^ + *b*𝔼 [*k*^2^] *δ*(*t* − *s*) d*t*d*s*. Consequently, using that *b*𝔼 [*k*] = *Kr*, we obtain:

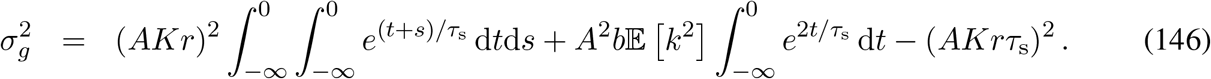

This implies that

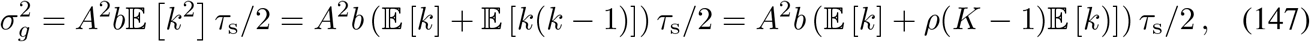

where the last equality follows from definition (140). Using once more that *b*𝔼 [*k*] = *Kr*, we have

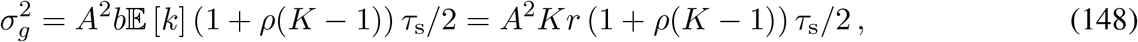

which shows that spiking correlations impact conductance variability whenever *ρ* is large or of order 1*/K*. This also shows that in the presence of instantaneous input synchrony, the amplitude-independent quantity *Q* becomes correlation-independent:

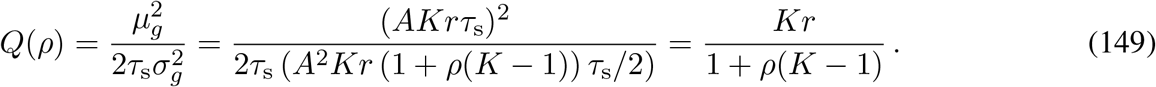

#### 3.3 Input drive with temporally-structured synchrony

We simulate temporally-structured synchrony by considering a doubly-stochastic model for synaptic input drive. In this model, the activation of individual synapses follow independent inhomogeneous Poisson processes with common rate given as a stationary CIR process *Z* satisfying:

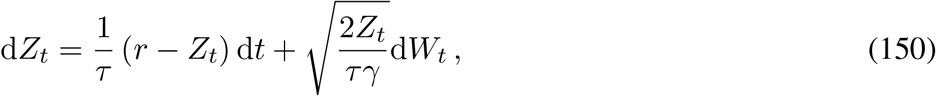

As before, the conductance process resulting from a doubly-stochastic synaptic-input model is specified as the inhomogeneous shot noise

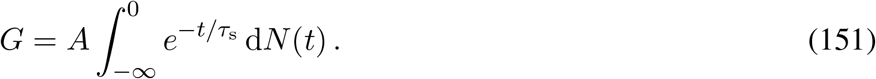

where the overall inhomogeneous Poisson process *N* has rate *KZ*(*t*). The mean stationary conductance can be computed as

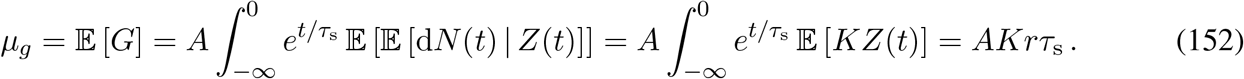

The stationary conductance variance can be evaluated as

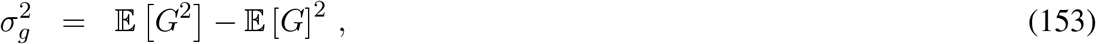

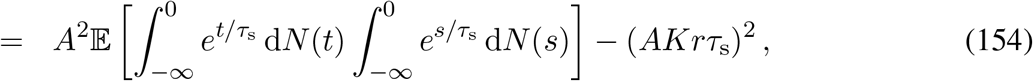

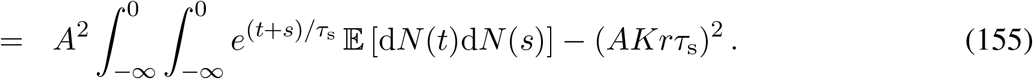

For a doubly-stochastic processes with common rate *Z*, we have

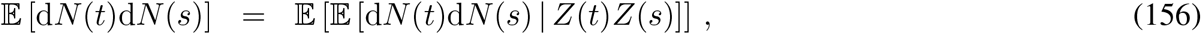

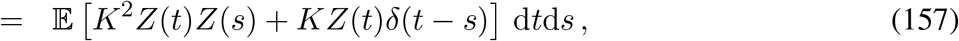

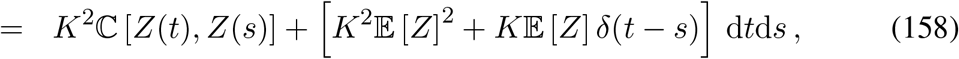

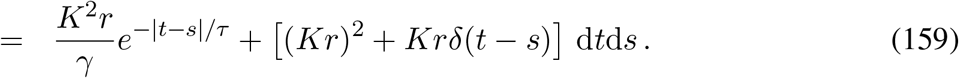

Injecting the above relation in expression (155) yields

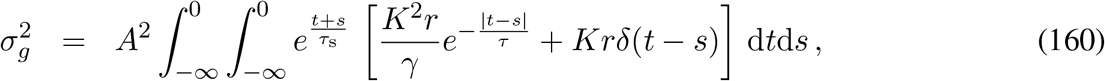

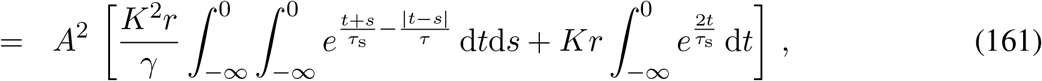

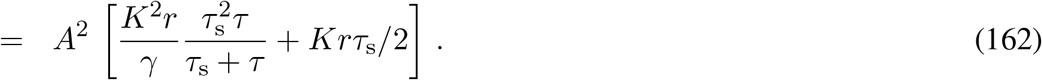

Thus, we obtain the amplitude-independent quantity as a function of the correlation time scale *τ* as

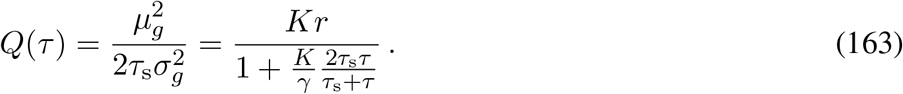

Suppose one measure a biophysically realistic correlation coefficient of *ρ*(∞) = 0.1 in large time bins *T*→ ∞. Then one deduce from… that *γ/*(2*τ*) ≃10. At the same time, we have *τ*_*s*_*/*(*τ* + *τ*_*s*_) ≃1*/*3 so that on can estimate:

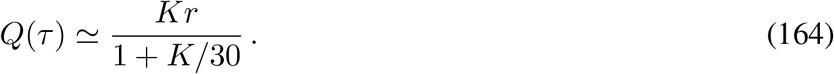

showing that even for moderately large number of synaptic contacts, *Q* can drastically underestimate the overall input rate *Kr* in the presence of temporally-structured synchrony.

## Bibliography

1. D. Tolhurst, J. A. Movshon, I. Thompson, The dependence of response amplitude and variance of cat visual cortical neurones on stimulus contrast. Experimental brain research 41, 414–419 (1981).

2. P. Kara, P. Reinagel, R. C. Reid, Low response variability in simultaneously recorded retinal, thalamic, and cortical neurons. Neuron 27, 635–646 (2000).

3. M. Carandini, Amplification of Trial-to-Trial Response Variability by Neurons in Visual Cortex. PLoS Biol 2, E264 (2004).

4. M. M. Churchland et al., Stimulus onset quenches neural variability: a widespread cortical phenomenon. Nat Neurosci 13, 369–378 (2010).

5. C. van Vreeswijk, H. Sompolinsky, Chaos in neuronal networks with balanced excitatory and inhibitory activity. Science 274, 1724–1726 (1996).

6. M. N. Shadlen, W. T. Newsome, The Variable Discharge of Cortical Neurons: Implications for Connectivity, Computation, and Information Coding. The Journal of Neuroscience 18, 3870–3896 (1998).

7. C. F. Stevens, A. M. Zador, Input synchrony and the irregular firing of cortical neurons. Nature Neuroscience 1, 210–217 (1998).

8. T. P. Vogels, L. F. Abbott, Signal Propagation and Logic Gating in Networks of Integrate- and-Fire Neurons. The Journal of Neuroscience 25, 10786–10795 (2005).

9. A. Bell, Z. F. Mainen, M. Tsodyks, T. J. Sejnowski, “Balancing” of conductances may explain irregular cortical spiking. La Jolla, CA: Institute for Neural Computation Technical Report INC-9502, (1995).

10. E. Schneidman, B. Freedman, I. Segev, Ion Channel Stochasticity May Be Critical in Determining the Reliability and Precision of Spike Timing. Neural Computation 10, 1679–1703 (1998).

11. G. Hennequin, Y. Ahmadian, D. B. Rubin, M. Lengyel, K. D. Miller, The Dynamical Regime of Sensory Cortex: Stable Dynamics around a Single Stimulus-Tuned Attractor Account for Patterns of Noise Variability. Neuron 98, 846–860 e845 (2018).

12. Z. Mainen, T. Sejnowski, Reliability of spike timing in neocortical neurons. Science 268, 1503–1506 (1995).

13. W. R. Softky, C. Koch, Cortical Cells Should Fire Regularly, But Do Not. Neural Computation 4, 643–646 (1992).

14. W. Softky, C. Koch, The highly irregular firing of cortical cells is inconsistent with temporal integration of random EPSPs. The Journal of Neuroscience 13, 334–350 (1993).

15. C. van Vreeswijk, H. Sompolinsky, Chaotic balanced state in a model of cortical circuits. Neural Comput 10, 1321–1371 (1998).

16. A. Renart et al., The asynchronous state in cortical circuits. Science 327, 587–590 (2010).

17. M. N. Shadlen, W. T. Newsome, Noise, neural codes and cortical organization. Current Opinion in Neurobiology 4, 569–579 (1994).

18. M. Tsodyks, T. J. Sejnowski, Rapid state switching in balanced cortical network models. Network: Computation in Neural Systems 6, (1995).

19. A. Y. Tan, B. D. Brown, B. Scholl, D. Mohanty, N. J. Priebe, Orientation selectivity of synaptic input to neurons in mouse and cat primary visual cortex. J Neurosci 31, 12339–12350 (2011).

20. M. C. Teich, C. Heneghan, S. B. Lowen, T. Ozaki, E. Kaplan, Fractal character of the neural spike train in the visual system of the cat. J Opt Soc Am A Opt Image Sci Vis 14, 529–546 (1997).

21. M. M. Churchland et al., Stimulus onset quenches neural variability: a widespread cortical phenomenon. Nature Neuroscience 13, 369–378 (2010).

22. R. L. T. Goris, J. A. Movshon, E. P. Simoncelli, Partitioning neuronal variability. Nature Neuroscience 17, 858–865 (2014).

23. G. R. Holt, W. R. Softky, C. Koch, R. J. Douglas, Comparison of discharge variability in vitro and in vivo in cat visual cortex neurons. Journal of Neurophysiology 75, 1806–1814 (1996).

24. M. Wehr, A. M. Zador, Balanced inhibition underlies tuning and sharpens spike timing in auditory cortex. Nature 426, 442–446 (2003).

25. L. I. Zhang, A. Y. Tan, C. E. Schreiner, M. M. Merzenich, Topography and synaptic shaping of direction selectivity in primary auditory cortex. Nature 424, 201–205 (2003).

26. N. J. Priebe, D. Ferster, Direction selectivity of excitation and inhibition in simple cells of the cat primary visual cortex. Neuron 45, 133–145 (2005).

27. B. H. Liu et al., Broad inhibition sharpens orientation selectivity by expanding input dynamic range in mouse simple cells. Neuron 71, 542–554 (2011).

28. A. S. Ecker et al., Decorrelated neuronal firing in cortical microcircuits. Science 327, 584–587 (2010).

29. Z.F. Kisvárday et al., Synaptic targets of HRP-filled layer III pyramidal cells in the cat striate cortex. Experimental Brain Research 64, 541–552 (1986).

30. B. Hellwig, A quantitative analysis of the local connectivity between pyramidal neurons in layers 2/3 of the rat visual cortex. Biological Cybernetics 82, 111–121 (2000).

31. T. Binzegger, R. J. Douglas, K. A. Martin, A quantitative map of the circuit of cat primary visual cortex. J Neurosci 24, 8441–8453 (2004).

32. H. Markram et al., Reconstruction and Simulation of Neocortical Microcircuitry. Cell 163, 456–492 (2015).

33. L. A. Becker, B. Li, N. J. Priebe, E. Seidemann, T. Taillefumier, Exact Analysis of the Subthreshold Variability for Conductance-Based Neuronal Models with Synchronous Synaptic Inputs. Phys Rev X 14, (2024).

34. J. C. Anderson, R. J. Douglas, K. A. Martin, J. C. Nelson, Map of the synapses formed with the dendrites of spiny stellate neurons of cat visual cortex. J Comp Neurol 341, 25–38 (1994).

35. R. J. Douglas, C. Koch, M. Mahowald, K. A. Martin, H. H. Suarez, Recurrent excitation in neocortical circuits. Science 269, 981–985 (1995).

36. S. Shinomoto, Y. Miyazaki, H. Tamura, I. Fujita, Regional and laminar differences in in vivo firing patterns of primate cortical neurons. J Neurophysiol 94, 567–575 (2005).

37. S. J. Kuhlman et al., A disinhibitory microcircuit initiates critical-period plasticity in the visual cortex. Nature 501, 543–546 (2013).

38. G. A. Wildenberg et al., Primate neuronal connections are sparse in cortex as compared to mouse. Cell Rep 36, 109709 (2021).

39. S. Loomba et al., Connectomic comparison of mouse and human cortex. Science 377, eabo0924 (2022).

40. M. R. Cohen, A. Kohn, Measuring and interpreting neuronal correlations. Nature Neuroscience 14, 811–819 (2011).

41. E. Schneidman, M. J. Berry, 2nd, R. Segev, W. Bialek, Weak pairwise correlations imply strongly correlated network states in a neural population. Nature 440, 1007–1012 (2006).

42. D. Golomb, J. Rinzel, Clustering in globally coupled inhibitory neurons. Physica D: Nonlinear Phenomena 72, 259–282 (1994).

43. D. Hansel, H. Sompolinsky, Chaos and synchrony in a model of a hypercolumn in visual cortex. J. Comp. Neurosci. 3, 7–34 (1996).

44. Y. Zerlaut, S. Zucca, S. Panzeri, T. Fellin, The Spectrum of Asynchronous Dynamics in Spiking Networks as a Model for the Diversity of Non-rhythmic Waking States in the Neocortex. Cell Reports 27, 1119-1132.e1117 (2019).

45. W. Bair, E. Zohary, W. T. Newsome, Correlated firing in macaque visual area MT: time scales and relationship to behavior. J Neurosci 21, 1676–1697 (2001).

46. M. A. Smith, A. Kohn, Spatial and Temporal Scales of Neuronal Correlation in Primary Visual Cortex. Journal of Neuroscience 28, 12591–12603 (2008).

47. J. F. Mitchell, K. A. Sundberg, J. H. Reynolds, Spatial attention decorrelates intrinsic activity fluctuations in macaque area V4. Neuron 63, 879–888 (2009).

48. M. Okun et al., Diverse coupling of neurons to populations in sensory cortex. Nature 521, 511–515 (2015).

49. T. A. Engel et al., Selective modulation of cortical state during spatial attention. Science 354, 1140–1144 (2016).

50. R. Zeraati et al., Intrinsic timescales in the visual cortex change with selective attention and reflect spatial connectivity. Nat Commun 14, 1858 (2023).

51. A. Kohn, M. A. Smith, Stimulus Dependence of Neuronal Correlation in Primary Visual Cortex of the Macaque. Journal of Neuroscience 25, 3661–3673 (2005).

52. R. Zeraati, T. A. Engel, A. Levina, A flexible Bayesian framework for unbiased estimation of timescales. Nature Computational Science 2, 193–204 (2022).

53. R. L. T. Goris, C. M. Ziemba, J. A. Movshon, E. P. Simoncelli, Slow gain fluctuations limit benefits of temporal integration in visual cortex. Journal of Vision 18, 8 (2018).

54. X. J. Wang, G. Buzsaki, Gamma oscillation by synaptic inhibition in a hippocampal interneuronal network model. J Neurosci 16, 6402–6413 (1996).

55. M. Pospischil et al., Minimal Hodgkin–Huxley type models for different classes of cortical and thalamic neurons. Biological Cybernetics 99, 427–441 (2008).

56. K. Padmanabhan, N. N. Urban, Intrinsic biophysical diversity decorrelates neuronal firing while increasing information content. Nat Neurosci 13, 1276–1282 (2010).

57. D. H. Hubel, T. N. Wiesel, Receptive fields, binocular interaction and functional architecture in the cat’s visual cortex. J. Physiol. (Lond.) 160, 106–154 (1962).

58. W. H. Bosking, Y. Zhang, B. Schofield, D. Fitzpatrick, Orientation selectivity and the arrangement of horizontal connections in tree shrew striate cortex. J Neurosci 17, 2112–2127 (1997).

59. H. Ko et al., Functional specificity of local synaptic connections in neocortical networks. Nature 473, 87–91 (2011).

60. L. Cossell et al., Functional organization of excitatory synaptic strength in primary visual cortex. Nature 518, 399–403 (2015).

61. B. Scholl, D. E. Wilson, D. Fitzpatrick, Local Order within Global Disorder: Synaptic Architecture of Visual Space. Neuron 96, 1127 - 1138.e1124 (2017).

62. A. Litwin-Kumar, B. Doiron, Slow dynamics and high variability in balanced cortical networks with clustered connections. Nature Neuroscience 15, 1498–1505 (2012).

63. R. Darshan, W. E. Wood, S. Peters, A. Leblois, D. Hansel, A canonical neural mechanism for behavioral variability. Nature Communications 8, 15415 (2017).

64. R. Rosenbaum, M. A. Smith, A. Kohn, J. E. Rubin, B. Doiron, The spatial structure of correlated neuronal variability. Nature Neuroscience 20, 107–114 (2017).

65. N. Brunel, V. Hakim, Fast global oscillations in networks of integrate-and-fire neurons with low firing rates. Neural Comput 11, 1621–1671 (1999).

66. C. van Vreeswijk, F. Farkhooi, Fredholm theory for the mean first-passage time of integrate-and-fire oscillators with colored noise input. Phys Rev E 100, 060402 (2019).

67. A. Sanzeni, M. H. Histed, N. Brunel, Emergence of Irregular Activity in Networks of Strongly Coupled Conductance-Based Neurons. Phys. Rev. X 12, 011044 (2022).

68. J. J. Pattadkal, B. V. Zemelman, I. Fiete, N. J. Priebe, Primate neocortex performs balanced sensory amplification. Neuron 112, 661-675.e667 (2024).

69. M. Pachitariu, S. Sridhar, J. Pennington, C. Stringer, Spike sorting with Kilosort4. Nat Methods 21, 914–921 (2024).

70. J. H. Siegle et al., Survey of spiking in the mouse visual system reveals functional hierarchy. Nature 592, 86–92 (2021).

71. M. L. Hines, N. T. Carnevale, The NEURON simulation environment. Neural Comput 9, 1179–1209 (1997).

72. R. A. McDougal et al., Twenty years of ModelDB and beyond: building essential modeling tools for the future of neuroscience. J Comput Neurosci 42, 1–10 (2017).

